# The E3 ubiquitin ligase Mind bomb 1 enhances nuclear import of viral DNA by inactivating a virion linchpin protein that suppresses exposure of virion pathogen-associated molecular patterns

**DOI:** 10.1101/2020.08.29.242826

**Authors:** Michael Bauer, Alfonso Gomez-Gonzalez, Maarit Suomalainen, Silvio Hemmi, Urs F. Greber

## Abstract

In eukaryotic cells, genomes from incoming DNA viruses mount two opposing reactions, viral gene expression and innate immune response, depending on genome exposure (uncoating) to either RNA-polymerases or DNA sensors. Here we show that adenovirus particles contain a tunable linchpin protein with a dual function: response to host cues for scheduled DNA release into the nucleus, and innate immunity suppression by preventing unscheduled DNA release. Scheduled DNA release required the proteasome and ubiquitination of the viral core protein V. Cells lacking the E3 ligase Mind bomb 1 (Mib1) were resistant to wild-type adenovirus infection. Viruses lacking protein V or bearing non-ubiquitinable protein V, however, readily infected Mib1 knockout cells, yet were less infectious than wild-type virus. Their genomes were poorly imported into the nucleus and remained uncoated in the cytosol, thereby enhancing chemokine and interferon production through the DNA sensor cGAS. Our data uncover how the ubiquitin-proteasome system controls the function of a virion linchpin protein suppressing pathogen-associated molecular patterns and triggers viral DNA uncoating at the nuclear pore complex for nuclear import and infection.

## INTRODUCTION

Pathogen-associated molecular patterns (PAMPs) activate the immune system as early as pathogens enter a cell, which facilitates adaptive immunity and coordinated cell defense ^1, 2^. PAMPs comprise a range of features, including DNA, double-stranded RNA, lipopolysaccharides or cytosolic glycans. They are decoded by pattern recognition receptors (PRRs), for example Toll-like receptors, DNA- and RNA-sensors such as cyclic GMP-AMP synthase (cGAS), nucleotide-binding oligomerization domain (NOD)-like receptors (NLRs), retinoid acid-inducible gene I (RIG-I)-like receptors (RLRs), or cytosolic lectins, which can lead to inflammatory responses ^3, 4, 5, 6^. The innate immune response is limited in extent and duration by both self-regulation and by the intruding pathogen. The work here uncovers that double-stranded DNA-PAMPs from incoming adenovirus (AdV) particles are effectively sequestered away from cytosolic sensors by the action of a virion protein functioning as a linchpin that secures the integrity of the incoming particles. This security mechanism prevents a massive cytokine response induced by unscheduled disassembly of the incoming capsid in the cytoplasm and exposure of the viral genome to the DNA sensor cGAS. The function of this linchpin can be relieved by targeted ubiquitination of protein V and allows viral genome release at the nuclear pore complex (NPC) for nuclear import and infection.

Ubiquitination tunes a wide range of cellular processes, including DNA-damage response, protein trafficking or autophagy, and generally results in proteasomal degradation or signal transduction ^7^. Ubiquitination starts with the linkage of a single ubiquitin peptide 76 amino acids in length to the ε-amino group of a lysine residue (K) via the C-terminal glycine (G) of ubiquitin ^8, 9^. This process comprises three distinct catalysts, E1 ubiquitin activation, E2 ubiquitin conjugation and E3 ubiquitin ligation enzymes ^9^. The latter control the substrate specificity in the ubiquitin transfer reaction and define the type of ubiquitin linkage ^10^. Proteins with ubiquitin binding domains recognize ubiquitinated proteins, and specialized ubiquitin hydrolases remove ubiquitin ^11^. Ubiquitin itself can be modified with ubiquitin by isopetide linkage at one of its lysine residues or at the N-terminal methionine residue, which is the basis of ubiquitin chain signaling through large chemical diversity, and affects almost any protein in a mammalian cell. Branched poly-ubiquitins with K11- or K48-linked chains appear to be particularly good substrates for binding to the proteasome and triggering substrate degradation ^12^.

Viruses are perhaps best known to take advantage of the ubiquitin-proteasome system (UPS) to ubiquitinate and degrade host factors that restrict their replication, such as the tumor suppressor protein p53 targeted by the HPV E6, E7 proteins, the AdV E1B-55K/E4Orf6 complex, or the HSV1 ICP0 protein reviewed in ^13^. Here we identified a novel role of the UPS in virus infection, namely to regulate virion stability and DNA genome uncoating in cells. Virion stability is critical for entry and egress, and allows viruses to deliver and express their genomes in infected cells, form progeny and spread to uninfected cells ^14^. Viruses bind to animal cells and trigger an entry process that uncoats their genetic information in a stepwise manner ^15, 16, 17^. Stepwise uncoating involves conformational flexibility and requires host cues, a universal feature of any virus particle ^18^. Conformational flexibility has been known as metastability, for example, the spring-loaded hemagglutinin glycoprotein of the enveloped influenza virus unloads upon low pH triggers ^19, 20^. At the level of nonenveloped viruses, conformational changes are triggered by receptor binding, alterations in chemical solutes, such as redox, calcium depletion or low pH in SV40, reovirus, polyomavirus or rhinovirus, respectively ^21, 22, 23, 24^, as well as mechanical cues ^25, 26, 27, 28^. Additional cues for uncoating are ubiquitination in the case of influenza virus, vaccinia virus, dengue virus, and AdV ^29, 30, 31, 32^. AdVs have a long history as vectors in gene therapy and vaccination, including coronavirus vaccine development, currently in late stages of clinical trials ^33, 34, 35, 36^. Human AdVs infect the gastrointestinal and respiratory tract as well as eyes ^37, 38, 39^. The viruses persist in a subset of cells following acute infection and reactivate during episodes of compromised immunity, e.g. after organ transplantation or anti-inflammatory treatment of autoimmune disease ^38, 40, 41^.

Here, we show that the RING-type E3 ubiquitin ligase Mind bomb 1 (Mib1) controls the stability of a critical AdV linchpin, protein V, by triggering its dissociation from the DNA core at the NPC, a process which also requires the activity of the proteasome. Mib1 catalyzes the transfer of polyubiquitin chains with several different linkages, including K48 ^42, 43, 44^, localizes to centriolar satellites ^43^, and is involved in developmental pathways, such as Notch signaling ^45^, ciliogenesis ^46^, and centriole biogenesis ^42^. It dynamically controls the sociology of the perinuclear zone, including virions of infected cells. Using Mib1 knock-out (KO) cells, mass spectrometry, site-directed mutagenesis, and cytokine profiling we demonstrate that the DNA core-associated protein V serves as a key stabilizer protecting cytoplasmic virions from premature (unscheduled) DNA-release. The work here provides crucial insight into the cell biology of DNA delivery into cell nuclei, a key step in viral disease, vaccination, and synthetic biology.

## RESULTS

### Proteasome activity is required for capsid disassembly

We described earlier that the enzymatic activity of the E3 ubiquitin ligase Mib1 triggers the uncoating of the adenoviral DNA from virions docked at the NPC ^32^. Here, we first determined whether the proteasome was involved in capsid disassembly and genome release. AdV-C5 infection in HeLa cells treated with two distinct proteasome inhibitors, MG132 and MLN9708, was strongly reduced 24 h post infection (pi) (**Fig. 1A**). Dose-response curves indicated inhibitory concentrations with 50% efficacy (IC_50_) of 0.176 and 0.082 μM, respectively. Very similar results were obtained in human diploid fibroblasts immortalized with telomerase (HDF-TERT) (**Fig. S1**). MLN9708 wash-out and wash-in experiments at 2 and 5 h pi, respectively, showed strong inhibition of infection, suggesting that viral entry as well as replication require an active proteasome (**Fig. 1B**). Using confocal fluorescence microscopy, we tracked viral genomes in HeLa cells inoculated with EdC-labeled AdV-C5 in the presence of MLN9708. MLN9078 strongly reduced the number of viral genomes released from the capsids, demonstrating that the proteasome is required for the release of the viral genome from incoming virions (**Fig. 1C, 1D**). To test if proteasome inhibition affected critical events upstream of DNA release, we allowed viral particles to reach the nucleus in Mib1-KO cells (sgMib1) for 3 h, and induced the expression of GFP-tagged Mib1 with doxycycline (**Fig. 1E**). In the absence of proteasomal inhibitors, DNA was readily released from the capsids, whereas the presence of MG132 strongly reduced the levels of capsid-free vDNA, indicating that proteasomal activity is required specifically for the release of vDNA from capsids docked at the NPC.

**Figure 1:**
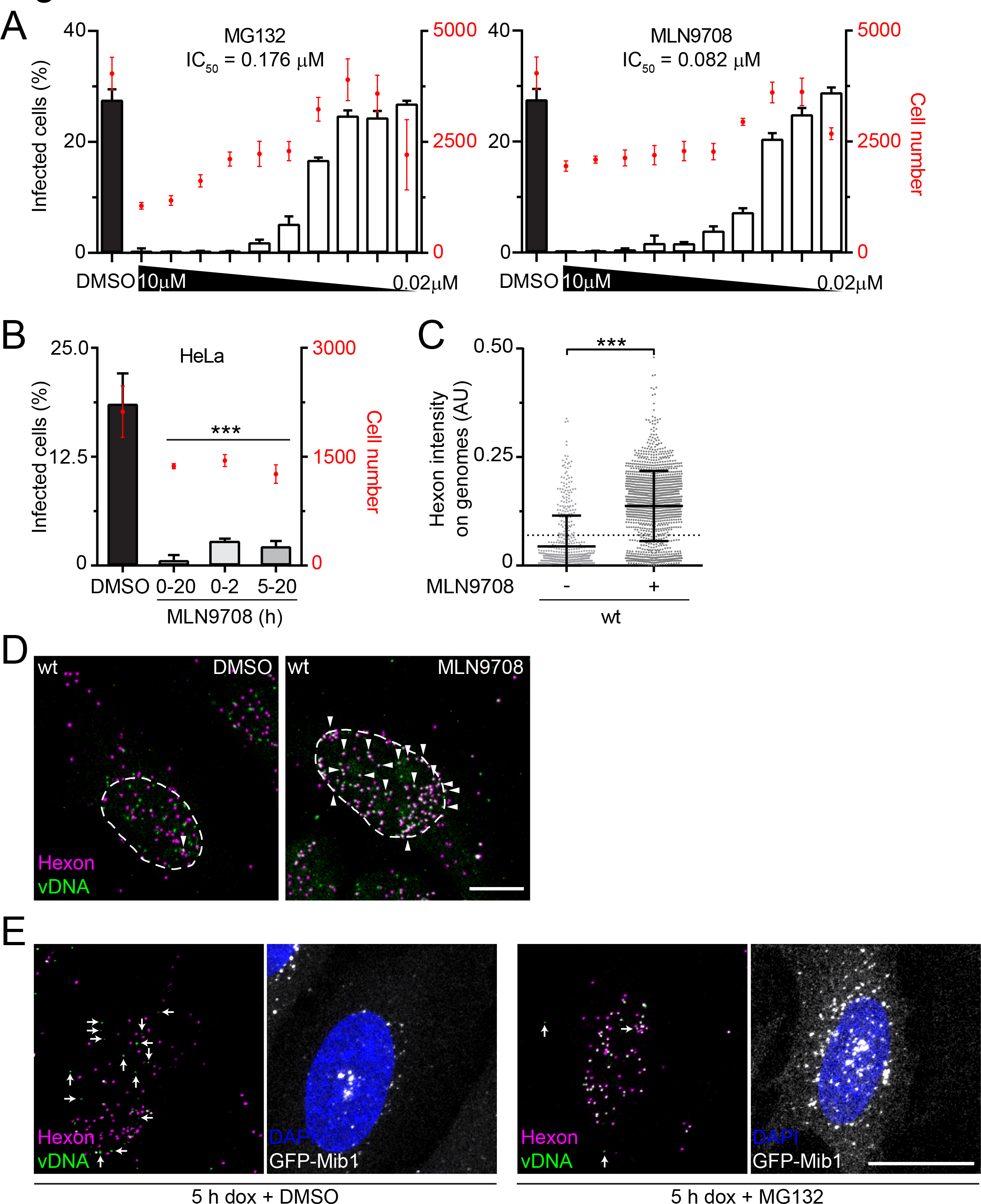
Proteasome activity is required for capsid disassembly at the NPC. (A) HeLa cells were infected with AdV-C5 at a MOI of 0.3 for 24 h in the presence of DMSO or varying concentrations of proteasome inhibitors MG132 or MLN9708. After fixation, cells were stained with anti-pVI and DAPI. Infection was scored by percentage of pVI-positive nuclei. Data are shown as mean ± SD. (B) HeLa cells were infected with AdV-C5 at a MOI of 0.3 for 20 h. MLN9708 (10 μM) was present for the entire infection (0-20), only during entry (0-2), or during the replication stages (5-20). Cells were fixed and processed as in (A). Data is shown as mean ± SD. Statistical significance was assessed using a one-way ANOVA with Holm-Sidak correction for multiple comparisons. ***, p<0.001. (C) HeLa cells were incubated with genome-labeled AdV-C5 for 30 min at 37 °C. The inoculum was replaced with fresh medium, and cells were fixed after 150 min. MLN9708 (10 μM) was present during the entire experiment. Incoming capsids were stained with anti-hexon, and viral genomes were visualized by click chemistry with an N3-AlexaFluor 488. Nuclei were stained with DAPI and cell outlines with succinimidyl ester. Cells were analyzed by confocal microscopy. Viral capsids were segmented and their corresponding vDNA intensity was plotted. Statistical significance was assessed by using a non-parametric ANOVA (Kruskal-Wallis test) with Dunn’s correction for multiple comparisons. ***, p<0.001. (D) Representative images from the dataset collected in (C). Images are maximum projections. Nuclear outlines are based on DAPI. Arrowheads indicate viral particles containing vDNA. Scale bar, 10 μm. (E) HeLa-sgMib1 cells carrying a tetracycline-inducible GFP-Mib1 cassette were incubated for 1 h with wt AdV-C5-EdC. Unbound virus was washed away, and internalized virus capsids were allowed to reach the nucleus for another 2 h, after which cells were treated with 1 μg/mL doxycycline plus DMSO or 10 μM MG132 for 5 h. Then cells were fixed, stained for vDNA, hexon, and nuclei, and imaged via confocal fluorescence microscopy. Images are maximum projections. Arrows indicate free viral genomes. Scale bar, 10 μm.

### Adenovirus core protein V is ubiquitinated during entry and removed from the virion at the NPC

Colocalization of EGFP-tagged Mib1 and viral capsids at the nuclear envelope occurs frequently as shown before ^32^. We used an affinity purification mass spectrometry (MS) based strategy to test the hypothesis that a viral capsid protein serves as a ubiquitination substrate of Mib1. HeLa cells were infected with AdV-C5 at high multiplicity of infection (MOI) for 2 h, lysed under strong denaturing conditions, and digested with trypsin protease. Trypsin cleaves the polypeptide chain C-terminal of K and arginine (R) residues, and leaves a di-glycine (Gly, G) remnant on the ε-amino group of ubiquitinated K residues, thus creating a peptide motif specifically recognized by a monoclonal antibody ^47, 48^. Immunoprecipitates of ubiquitinated cellular and viral peptides analyzed by liquid chromatography tandem MS (LC-MS/MS) contained peptides from over 5000 cellular proteins, as well as four viral proteins: penton base, IIIa, V, and VI (**Fig. 2A, 2B, 2C**). The virion proteins were not previously described to be ubiquitinated, except for protein VI ^49^. Unlike penton base, IIIa and VI ^26, 50^, a large fraction of protein V remains inside the virus particle until capsid disassembly at the NPC ^51^. We therefore investigated the role of protein V and the ubiquitination of K178 and K188 during the virus entry, and explored the dynamics of GFP-tagged protein V in incoming AdV-C2-GFP-V particles. Atto647-labeled AdV-C2-GFP-V particles (bearing a red fluorophore Atto647 and a green GFP-V) reached the nucleus of mScarlet-Mib1 expressing Mib1-KO cells (HeLa-sgMib1) some 30 min pi. Approximately 18 min after docking at the NPC, GFP-V puncta were rapidly discharged from the capsids (**Movie 1, Fig. 2D**). Of note, we never observed translocation of protein V puncta into the nucleus, while vDNA is either imported into the nucleus or misdelivered to the cytosol ^32, 52, 53^. GFP-V puncta were found to dissociate from capsids in control HeLa-sgNT, but not in HeLa-sgMib1 cells (**Fig. 2E**). It is therefore likely that the sudden loss of protein V from capsids at the NPC represents the final capsid disassembly step and accompanies the viral genome release.

**Figure 2:**
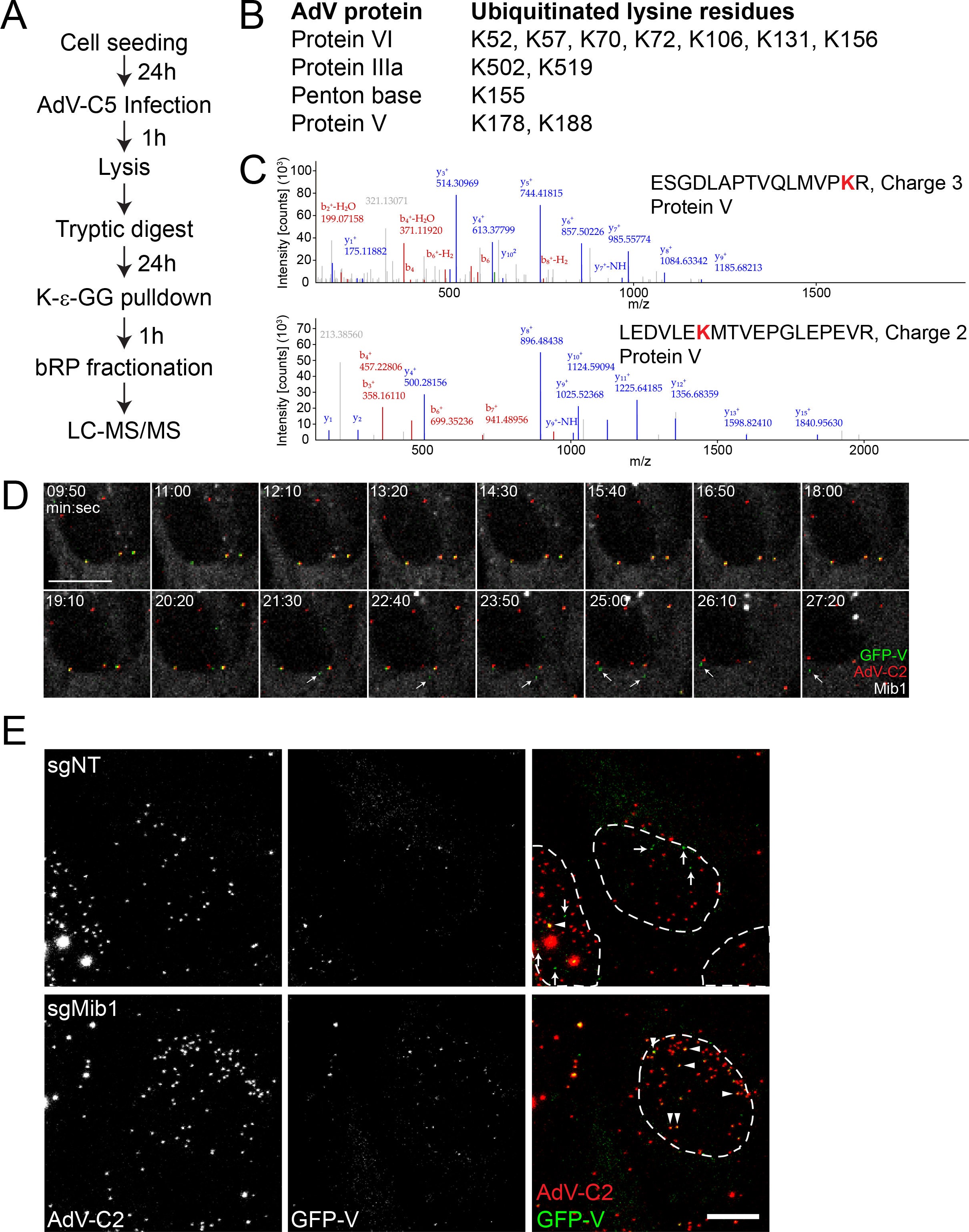
The core protein V is ubiquitinated during virus entry into cells and dissociates from the virion at the NPC. (A) Workflow of di-glycine immunoprecipitation and analysis by LC-MS/MS. (B) List of AdV-C5 proteins for which at least one peptide containing K-ε-GG residues were found. The corresponding Lys positions in the proteins are indicated. (C) Protein V ubiquitination sites were identified by mass spectrometry following protein digestion with LysC and trypsin, IP with di-glycine antibody, and LC MS/MS analysis. Two protein V ubiquitination sites were detected. Top, MS2 fragmentation spectra of the ESGDLAPTVQLMVPK(diGly)R peptide containing K178. Bottom, MS2 fragmentation spectra of peptide LEDVLEK(diGly)MTVEPGLEPEVR containing the K188 ubiquitination site. All matching b and y ions are annotated with their measured m/z (mass to charge) values. (D) Montage of selected timepoints from Movie S1. HeLa-sgMib1 cells expressing mScarlet-Mib1 were incubated with AdV-C2-GFP-V-atto647 for 30 min at 37 °C. The virus inoculum was removed, and cells were placed in a confocal spinning-disk microscope. Timestamps are in min:sec. Arrows indicate free GFP-V puncta after capsid dissociation. Three capsids containing GFP-V are visible at the nucleus, of which the two particles on the left discharge their GFP-V. Scale bar, 10 μm. (E) HeLa-sgNT or HeLa-sgMib1 cells were incubated with AdV-C2-GFP-V-atto647 for 30 min at 37 °C, after which the virus inoculum was removed, and cells were incubated for another 120 min. Cells were fixed, nuclei were stained with DAPI, and samples were imaged on a confocal laserscanning microscope. Images are maximum projections. Nuclear outlines are based on DAPI signal. Arrows indicate free GFP-V puncta, arrowheads show examples of virus particles positive for GFP-V. Scale bar, 10 μm.

### Virions lacking protein V are less dependent on Mib1 for infection and release vDNA prematurely

To determine how protein V affects the entry of the virion into host cells, we generated an AdV-C5 mutant deleted of the protein V coding region (AdV-C5-ΔV) (**Fig. 3A**). Particles isolated by cesium chloride gradient centrifugation and analyzed by SDS-PAGE and QuickBlue staining showed lack of protein V but no differences in the major virion proteins, including hexon, penton base, IIIa, fiber (IV), VI, and VII (**Fig. 3B**). Proteins VIII, IX, X, IVa2 and protease were not visualized in these gels. Sanger sequencing confirmed that the protein V coding region was removed from vDNA (**Fig. S2A**). Full analyses of the viral genome by next-generation sequencing revealed no other mutations in the viral genome (**Fig. S2B, and supporting material**). The integrity of the virions was further demonstrated by transmission electron microscopy of negatively stained specimens (**Fig. 3C**).

**Figure 3:**
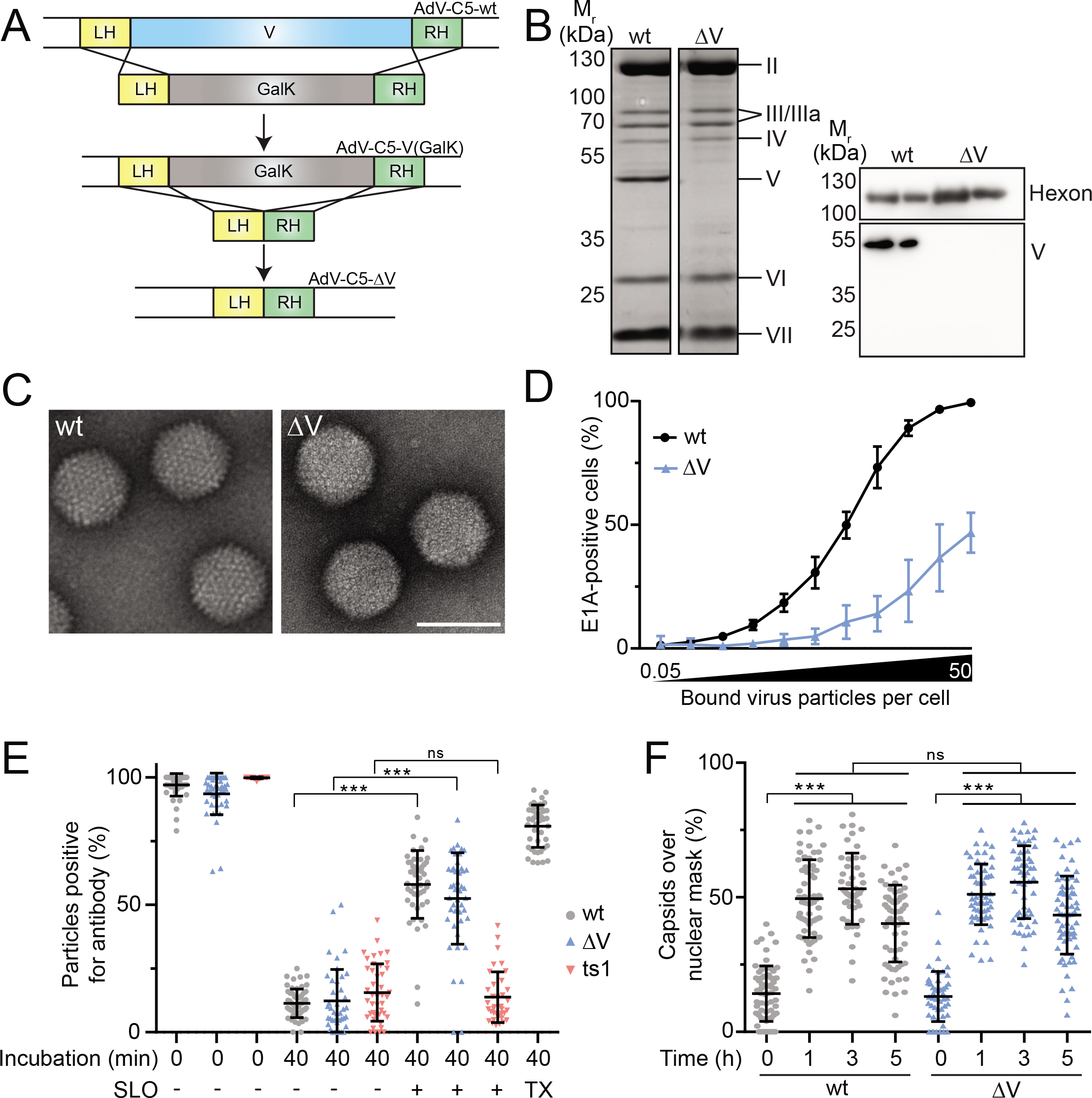
Viral particles lacking protein V are less infectious than the wild-type. (A) Recombineering strategy for the construction of AdV-C5-ΔV. Left homology (LH) and right homology (RH) arms were 45 bp long directly up- or downstream of the protein V coding region. In the first reaction, the protein V region was replaced with a GalK (galactokinase) cassette. In the second reaction, the GalK cassette was removed. The mutated AdV genome was then excised and transfected into 911 cells. Rescued virus was plaque-purified, expanded, and purified. (B) Purified AdV-C5 or AdV-C5-ΔV particles were lysed and loaded onto a gel. After SDS-PAGE, proteins in the gel were subjected to QuickBlue staining (left) or transferred to a PVDF membrane (right). Viral proteins Hexon and protein V were immunostained. Apparent molecular weights and corresponding viral proteins are indicated. (C) Electron micrographs of negatively stained AdV-C5 or AdV-C5-ΔV particles. Scale bar, 100 nm. (D) HeLa cells were incubated with AdV-C5 or AdV-C5-ΔV in the range of 0.05 to 50 bound viral particles per cell for 1 h on ice. The virus inoculum was replaced by fresh medium and cells were incubated at 37 °C for 20 h. After fixation, cells were stained with anti-E1A antibody and DAPI. Infection was scored by percentage of E1A-positive nuclei. Data are shown as mean ± SD. (E) A549 cells were incubated with atto565-labeled AdV-C5 or AdV-C5-ΔV for 1 h on ice. AlexaFluor 488-labeled AdV-C2 ts1 was used as a control. Virus inoculum was replaced by fresh medium and cells were either kept on ice or incubated at 37 °C for 40 min. Cell were permeabilized with Streptolysin-O and incubated with anti-Hexon or anti-Alexa488 antibodies. After fixation, cells were stained with the secondary antibodies, DAPI and succinimidyl ester. Virus particles were segmented based on their capsid label signal and classified based on their antibody signal. A Triton X-100 control was used to address the accessibility of the antibodies. Statistical significance was assessed by using a non-parametric ANOVA (Kruskal-Wallis test) with Dunn’s correction for multiple comparisons. ***, p<0.001; ns, non-significant. (F) A549 cells were incubated with AdV-C5 or AdV-C5-ΔV for 1 h on ice. Virus inoculum was replaced, and cells were either fixed or incubated at 37 °C for 1, 3, or 5 h. After fixation, cells were stained with anti-Hexon, DAPI, and succinimidyl ester. Virus particles were segmented and masked with the nucleus. Statistical significance was assessed as in (E).

Protein V promotes the assembly of newly formed viral particles ^54^, and AdV particles lacking protein V have been reported to form smaller plaques than wt AdV-C5, with the caveat that this old deletion mutant had many other mutations besides the deletion of V ^55^. To test if protein V was involved in the viral entry, we performed a single-round infection assay with AdV-C5-ΔV and parental AdV-C5 wild-type and measured the immediate early viral protein E1A 20 h pi. Inocula were adjusted to the number of particles bound per cell, as described earlier ^56^. To reach a certain level of E1A expression, a significantly higher number of AdV-C5-ΔV particles was required per cell, as compared to wild-type, indicating that particles lacking protein V were less infectious than wild-type (**Fig. 3D**). To investigate entry more closely, we performed analyses of single fluorescent particles using confocal light microscopy. These experiments revealed no difference in viral escape from the endosomes or trafficking to the nucleus between AdV-C5-ΔV and wild-type particles (**Fig. 3E, 3F**). However, the analyses of incoming viral genomes tagged with EdC nucleosides ^32, 52^ revealed significant differences between AdV-C5-ΔV and wild-type infections. While nearly all genomes were released from both wild-type and AdV-C5-ΔV capsids 3 h pi (**Fig. 4A, 4B**), the majority of wild-type virus genomes were delivered to the nucleus, yet most of the AdV-C5-ΔV genomes were in the cytoplasm (**Fig. 4A, 4C**). This defect in nuclear import of AdV-C5-ΔV vDNA likely explains the low infectivity of these particles. We next tested whether the absence of the ubiquitination substrate protein V would affect the infection of Mib1-KO cells. Strikingly, infection of Mib1-KO cells with AdV-C5-ΔV but not AdV-C5-ΔIX was readily possible, while wild-type infection was blocked (**Fig. 4D**). A higher number of released vDNA free from capsids was found in Mib1-KO cells, indicating that Mib1 was not involved in releasing vDNA from the AdV-C5-ΔV particles (**Fig. 4E, 4F**).

**Figure 4:**
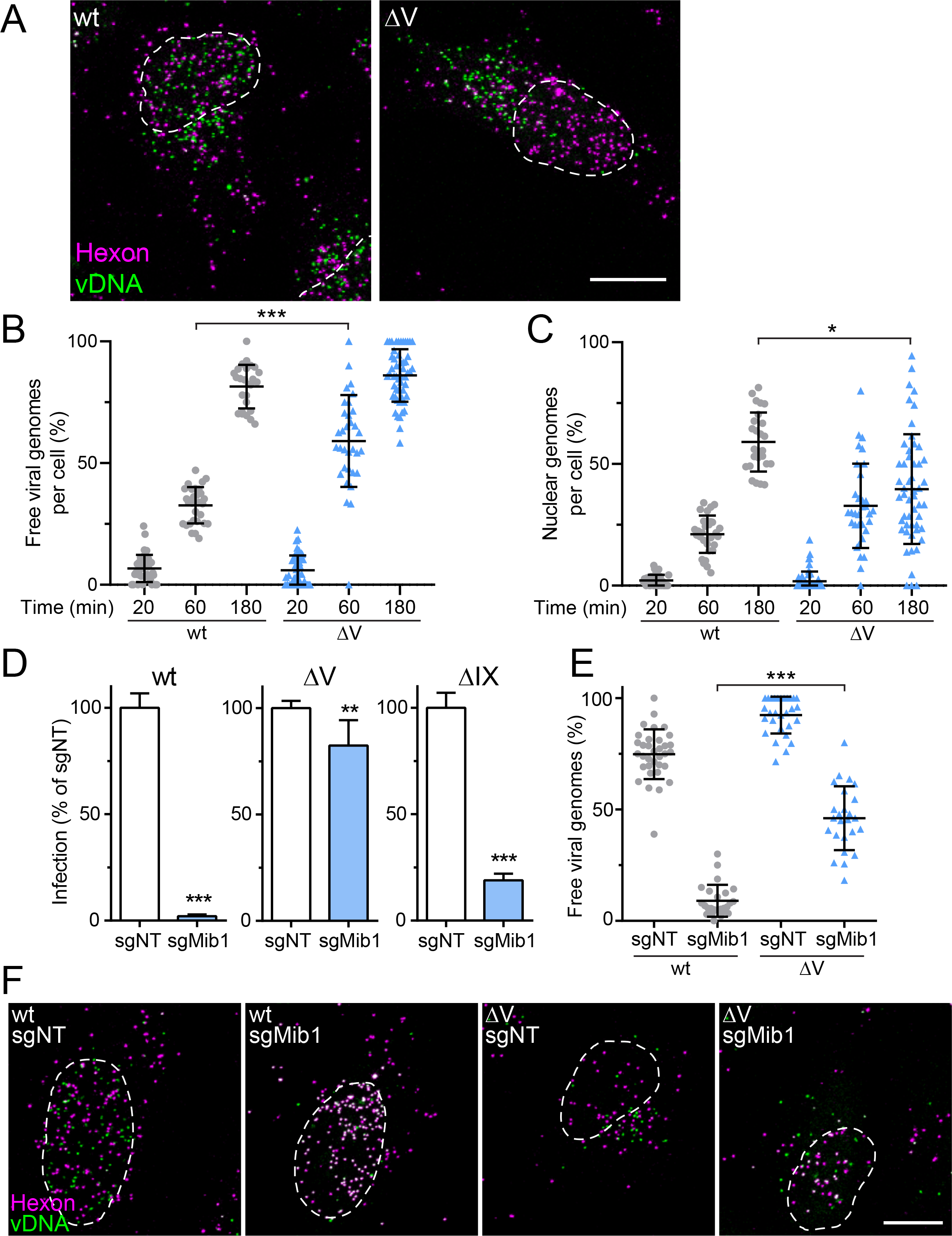
AdV-C5-ΔV particles deliver higher amounts of vDNA to the cytoplasm and are less dependent on Mib1 than wild-type AdV. (A) HeLa cells were incubated with genome-labeled AdV-C5 or AdV-C5-ΔV for 3 h. Cells were fixed and stained with anti-Hexon and vDNA using click chemistry. Nuclear outlines are based on the DAPI signal. Images are maximum projections. Scale bar, 10 μm. (B, C) HeLa cells were incubated with genome-labeled AdV-C5 or AdV-C5-ΔV for 20, 60, or 180 min and processed as in (A). Viral genomes were segmented and classified as free based on their Hexon intensity. Genomes over the nuclear mask were counted as nuclear. Each dot represents one cell. Data is shown as the mean ± SD. Statistical significance was assessed by using a nonparametric ANOVA (Kruskal-Wallis test) with Dunn’s correction for multiple comparisons. *, p<0.05; ***, p<0.001. (D) HeLa-sgNT and sgMib1 cells were infected with the indicated viruses at a MOI of 0.4. After 24 h, cells were fixed and stained with anti-pVI and DAPI. Nuclei were segmented based on DAPI signal and infection was based on pVI signal. Error bars represent mean ± SD. p values were assessed using unpaired t tests. **, p<0.01; ***, p<0.001. (E, F) HeLa-sgNT and sgMib1 cells were incubated with genome-labeled AdV-C5 or AdV-C5-ΔV for 3 h. Cells were fixed and stained with anti-Hexon and vDNA using click chemistry. Nuclear outlines are based on the DAPI signal. Images are maximum projections. Viral genomes were quantified as in (B). Statistical significance was assessed with a non-parametric ANOVA (Kruskal-Wallis test) with Dunn’s correction for multiple comparisons. ***, p<0.001. Scale bar, 10 μm.

### AdV-C5-ΔV are more thermo-sensitive than wild-type particles and release their genome in the cytoplasm before reaching the nucleus

The observation that AdV-C5-ΔV particles released their genome prematurely and did not require a ubiquitination cue from Mib1 suggested that they were less stable than wild-type particles. We assessed the thermal stability of the particles *in vitro* by measuring the fluorescence increase of the DNA intercalating dye DiYO-1, a highly sensitive end point assay for the disruption of the capsid shell ^51, 57^. At temperatures below 40 °C, both wild-type and AdV-C5-ΔV particles were largely intact, as indicated by a low level of DiYO-1 fluorescence, but AdV-C5-ΔV readily reached half maximal fluorescence at 42° C while wild-type virus remained intact up to about 47 °C (**Fig. 5A**). Accordingly, the infectivity of AdV-C5-ΔV particles was more thermo-sensitive than wild-type virus (**Fig. 5B**). This notion was reinforced by single virus particle analyses in cells, where more than 40% of the AdV-C5-ΔV particles lost their genome in the presence of the uncoating inhibitor leptomycin B (LMB) (**Fig. 5C, 5D**). Both wild-type and AdV-C5-ΔV infections were completely sensitive to LMB (**Fig. S3A**). LMB is a widely used nuclear export inhibitor ^58^, precludes the detachment of virions from microtubules near the nuclear envelope and thereby blocks virion attachment to the NPC and prevents nuclear import of viral DNA and infection ^59, 60^. The treatment of cells with protein degradation inhibitors, such as DBeQ blocking the p97 AAA-ATPase or MLN9708 against the proteasome, did not inhibit the premature release viral DNA from AdV-C5-ΔV (**Fig S3B, S3C**). Likewise, the depletion of microtubules with nocodazole had no effect (**Fig S3D, S3E**). Collectively, the results showed that the AdV-C5-ΔV particles were less stable than wild-type, and suggested that unscheduled release from AdV-C5-ΔV might be due to intrinsic particle instability in a crowded cytosol rather than a specific host anti-viral effect.

**Figure 5:**
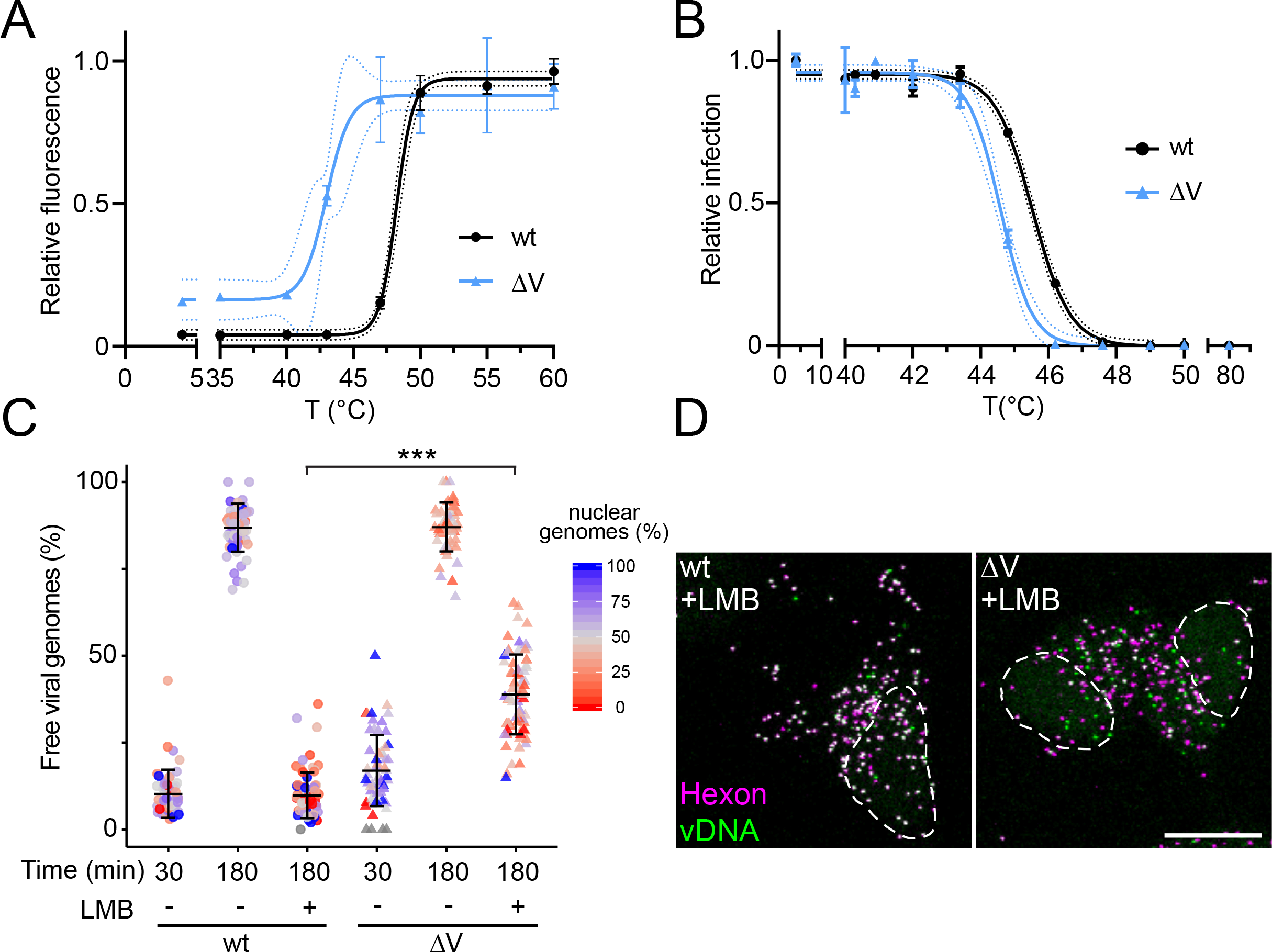
AdV-C5-ΔV particles are less thermo-stable and in the cytoplasm release vDNA more frequently than wild type. (A) Purified AdV-C5 or AdV-C5-ΔV particles were either kept at 4 °C or subjected to the indicated temperatures for 5 min. Samples were cooled down and incubated for 5 min with 5 μM DNA intercalating DiYO-1 before measuring their fluorescence. Fluorescence intensities were normalized to the maximum fluorescence at 60 °C. Data is shown as mean ± SD. (B) A549 cells were incubated with heat-treated AdV-C5 or AdV-C5-ΔV particles for 24 h. After fixation, cells were stained with anti-VI antibody and DAPI. Infection was scored by percentage of VI-positive nuclei relative to the non-treated samples. Data is shown as mean ± SD. (C, D) HeLa cells were incubated with genome-labeled AdV-C5 or AdV-C5-ΔV for 30 or 180 min in the absence or presence of 50 nM LMB. Cells were fixed and stained with anti-Hexon and vDNA using click chemistry. Data are shown as mean ± SD. Ratio of the number of capsid-free genomes over the nuclear mask has been color coded. Statistical significance was assessed by using a nonparametric ANOVA (Kruskal-Wallis test) with Dunn’s correction for multiple comparisons. ***, p<0.001. Nuclear outlines are based on the DAPI signal. Images are maximum projections. Scale bar, 10 μm.

### Protein V ubiquitination enhances AdV infectivity

The results so far suggested that the viral DNA core-associated protein V serves a ubiquitin responsive linchpin in the release of the viral DNA from the incoming capsid. We next assessed whether the ubiquitination of protein V at K178 and K188 was important for AdV infection, and created three AdV-C5 mutants, in which either one or both of these lysine residues was replaced by arginine, a basic amino acid that cannot serve as a ubiquitin acceptor site (**Fig. S4A**). HeLa-sgNT and sgMib1 cells infected with both single mutants and the double mutant (K178R, K188R, K178/188R) all showed strongly Mib1-dependent infections (**Fig. S4B**). Importantly, the doseresponse curves with the mutants were virtually identical to the wild-type, indicating that the mutant particles had the same specific infectivity as the wild-type (**Fig S4C**). We surmised that other lysines than K178 and K188 are ubiquitinated in the double mutant, and that these modifications were not detected by our MS analyses of protein V in the wild-type particles.

To abrogate any canonical ubiquitination of protein V, we changed all 26 lysine residues of protein V to arginines. Purified AdV-C5-V-KR particles incorporated the V-KR protein at comparable amounts as the parental AdV-C5 protein V, as shown by SDS-PAGE and QuickBlue protein staining (**Fig. 6A**). While the V-KR band appeared to migrate at a slightly faster than the wild-type protein V, there was no difference in any of the other capsid proteins resolved in the gel. Yet, the infectivity of AdV-C5-V-KR particles in HeLa cells was strongly impaired compared to the parental AdV-C5, comparable to the AdV-C5-ΔV mutant (**Fig. 6B**). The AdV-C5-V-KR particles released some of their vDNA but did not efficiently import this vDNA into the nucleus (**Fig. 6C, 6D**). In accordance, AdV-C5-V-KR infected HeLa-sgMib1 cells with greater efficiency than AdV-C5, demonstrating that this virus – similar to AdV-C5-ΔV – is less dependent on Mib1 for infection (**Fig. 6E**). The data showed that preventing ubiquitination at K178 and K188 had no effect on vDNA uncoating and infection, but replacing all lysine residues with arginine strongly reduced infection and dependence on Mib1.

**Figure 6:**
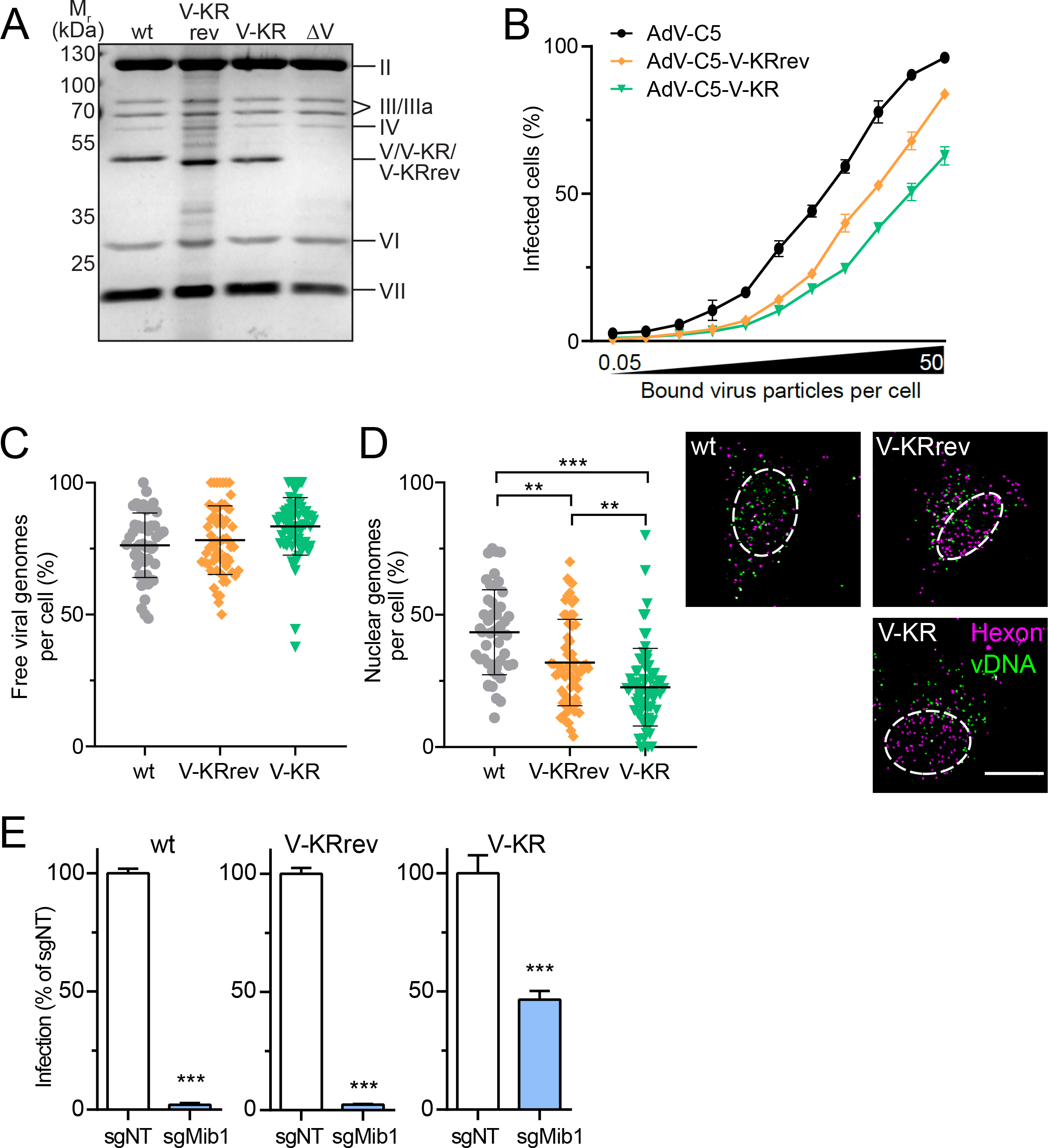
Protein V mutagenesis reveals that K178 and K188 are suffiicent for incoming AdV particles to respond to Mib1 infection cues. (A) Equal amounts of AdV-C5, AdV-C5-V-KRrev, AdV-C5-V-KR, and AdV-C5-ΔV were lysed and loaded onto a gel. After protein separation by SDS-PAGE, proteins were stained using QuickBlue. (B) HeLa cells were incubated with AdV-C5, AdV-C5-V-KRrev or AdV-C5-V-KR in the range of 0.05 to 50 bound viral particles per cell for 1 h on ice. The virus inoculum was replaced by fresh medium and the cells were incubated at 37 °C for 24 h. After fixation, cells were stained with anti-E1A and DAPI. Infection was scored by percentage of E1A-positive nuclei. Data is shown as mean ± SD. (C, D) HeLa cells were incubated with genome-labeled AdV-C5, AdV-C5-V-KRrev, or AdV-C5-V-KR for 30 min. Inocula were washed away, cells were incubated for a total of 180 min. Cells were fixed and stained for Hexon and vDNA using click chemistry. Nuclei were stained with DAPI. Data is shown as mean ± SD. Statistical significance was assessed using a non-parametric ANOVA (Kruskal-Wallis) with Dunn’s correction. Representative images show maximum projections. Nuclear outlines are based on DAPI signal. Scale bar, 10 μm. **, p<0.01; ***, p<0.001. (E) HeLa-sgNT and sgMib1 cells were infected with the indicated viruses at a MOI of 0.4. After 24 h, cells were fixed and stained with anti-pVI and DAPI. Nuclei were segmented based on DAPI signal and classified as infected based on their pVI signal. Data are shown as mean ± SD. p values were assessed using unpaired t tests. ***, p<0.001.

To test whether ubiquitination of Lys178 and Lys188 is sufficient to revert the phenotype of the V-KR mutant and restore the infection dependence on Mib1, we reverted the arginine residues to lysine at positions 178 and 188. The resulting mutant particles AdV-C5-V-KRrev efficiently incorporated the V-RKrev protein (**Fig 6A**). The AdV-C5-V-KRrev particles exhibited more efficient nuclear import of vDNA and higher infectivity than V-KR but were still less infectious than wild-type AdV-C5 (**Fig. 6B-D**). Nevertheless, AdV-C5-V-KRrev was strongly restricted in Mib1-KO cells, similar to AdV-C5 but unlike the AdV-C5-V-KR virus, suggesting that ubiquitination of protein V at K178 and K188 specifies the infection dependence on Mib1 (**Fig. 6E**).

### Strong induction of cytokines by AdV-C5-ΔV in a macrophage-like cell line

The results so far showed that tampering with the function of protein V reduces infectivity and increases the amounts of incoming vDNA in the cytoplasm. We next tested if the protein V mutants affected the induction of innate immune responses in the non-transformed, GM-CSF dependent, macrophage-like MPI-2 cell line ^61, 62^. Cells were infected with equal amounts of virus particles and mRNA levels of various cytokines were measured by reverse transcriptase quantitative polymerase chain reaction (RT-qPCR) at 5 and 10 h pi. Strikingly, virus particles lacking protein V led to higher levels of interleukin-1α (Il1a), Ccl2, Cxcl2, and Ccl5 mRNAs than wild-type (**Fig. 7A**). These results were confirmed by single cell analyses of the Ccl2 mRNA using RNA FISH assay with branched DNA signal amplification (**Fig. S5A**). Although V-KRrev and V-KR particles enhanced these cytokines compared to wild-type infection, their induction was not as pronounced as with AdV-C5-ΔV, in direct correlation with the levels of cytosolic vDNA. The results were corroborated by measuring type I IFN released from the infected MPI-2 cells. The assay used mouse embryonic fibroblasts expressing firefly luciferase under the control of the endogenous type I IFN inducible Mx2 promoter (MEF-Mx2-Luc-βKO) ^62^. Strikingly, infection with AdV-C5-ΔV led to a significantly higher IFN-β secretion than the wild-type virus. AdV-C5-V-KR and the V-KRrev increased IFN-β secretion as well, but not as prominently as AdV-C5-ΔV (**Fig. 7B**).

**Figure 7:**
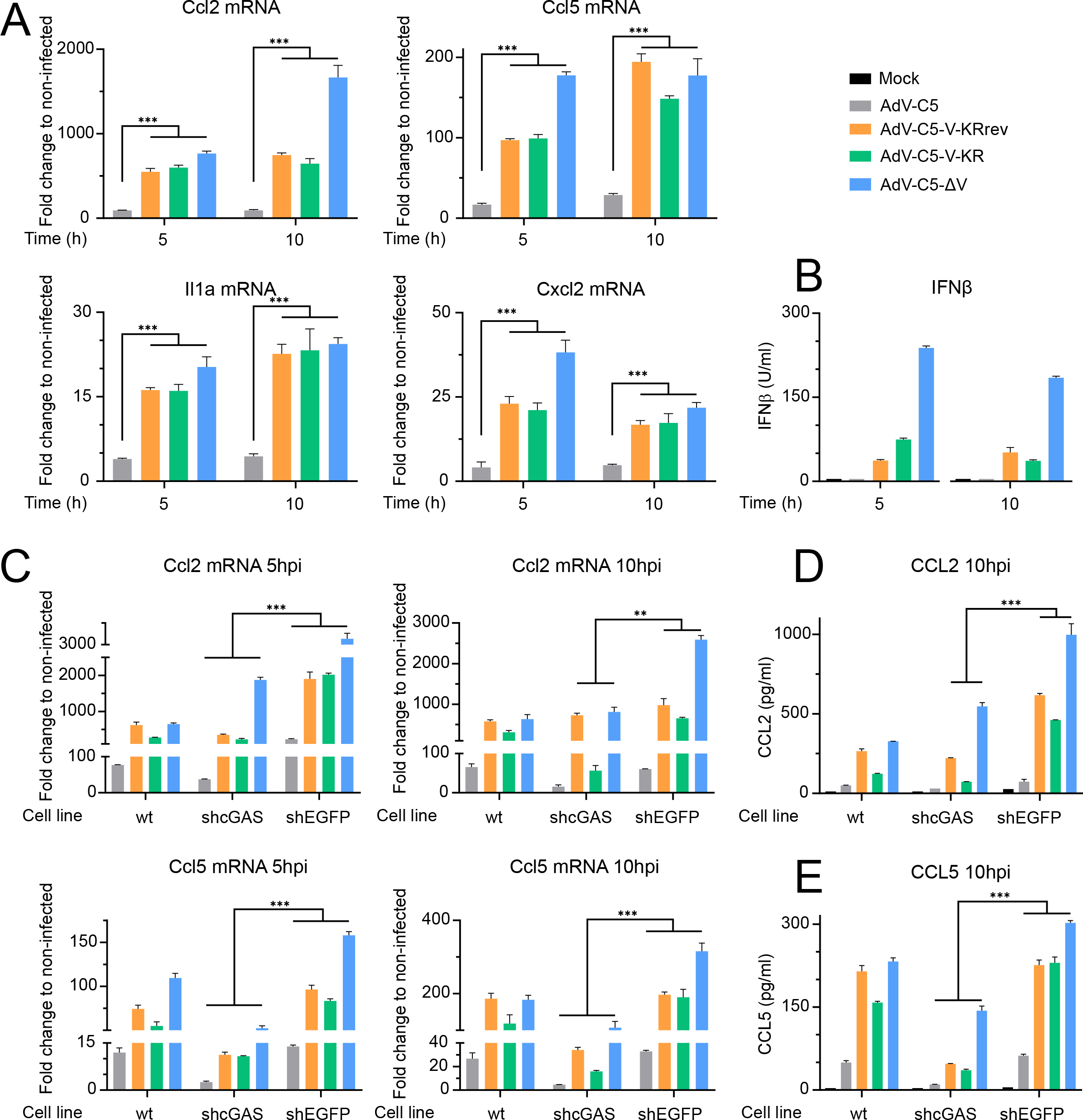
Protein V from incoming AdV tunes the induction of cytokines in a macrophage-like cell line early stages in infection. (A) MPI-2 cells were inoculated with AdV-C5, AdV-C5-VKRrev, AdV-C5-V-KR, or AdV-C5-ΔV in the range of 350 bound particles per cell for 30 min at 37 °C. Virus inoculum was replaced and cells were incubated at 37 °C for 5 or 10 h. RNA was extracted from the cellular fraction. Reverse-transcribed mRNA was amplified and measured through qPCR with primes for the indicated genes. Fold change in mRNA expression was addressed through the 2^-(ΔΔCt) method using HPRT cDNA as an endogenous control. Statistical significance was assessed by one-way ANOVA with Holm-Sidak correction for multiple comparisons. ***, p<0.001. (B) MEF-Mx2-Luc-βKO cells were inoculated with supernatants from MPI-2 wt, shcGAS, or shEGFP cells either mock or AdV infected. 20 h post inoculation cells were lysed and incubated with a luciferase substrate, followed by chemiluminescence measurement. (C) MPI-2 wt, shcGAS, or shEGFP cells were inoculated with AdV-C5, AdV-C5-VKRrev, AdV-C5-V-KR, or AdV-C5-ΔV as in (A). Samples were processed as described in (A). Statistical significance was assessed by one-way ANOVA with Holm-Sidak correction for multiple comparisons. ***, p<0.001. (D) Supernatant from MPI-2 wt, shcGAS or shEGFP cells mock or infected with AdV-C5, AdV-C5-VKRrev, AdV-C5-V-KR, or AdV-C5-ΔV were quantified with Sigma’s Mouse CCL2 ELISA Kit according to the manufacturer’s instructions. Statistical significance was assessed by one-way ANOVA with Holm-Sidak correction for multiple comparisons. ***, p<0.001. (E) Protein levels in the supernatant from MPI-2 wt, shcGAS or shEGFP cells mock-treated or infected with AdV-C5, AdV-C5-VKRrev, AdV-C5-V-KR, or AdV-C5-ΔV were quantified with a Mouse CCL5 ELISA Kit. Statistical significance was assessed by one-way ANOVA with Holm-Sidak correction for multiple comparisons. ***, p<0.001.

A major DNA-PAMP sensor and transducer in eukaryotic cells is the cGAS/STING branch ^6^, which detects also AdV DNA ^63, 64^. Upon DNA binding cGAS produces cGAMP which binds the adaptor protein STING and leads to the activation of TBK1. Active TBK1 phosphorylates IRF3 which then dimerizes, translocates to the nucleus, and promotes the expression of type I IFN. Significantly, the knockdown of cGAS by small hairpin (sh) RNA ^64^ reduced the induction of Ccl2 and Ccl5 mRNA in MPI-2 cells as compared to control shEGFP cells (**Fig. 7C and Fig. S5B**). The results were validated by ELISA assays showing that AdV-C5-ΔV increased the protein levels of CCL2 and CCL5 to much higher levels than the wild-type, notably in a cGAS-dependent manner (**Fig. 7D, 7E and Fig. S5**). Taken together, these results indicate that the presence of protein V in AdV particles reduced the levels of PAMPs in the cytoplasm and confined the innate response against the incoming virus (**Fig. 8**). Since protein V occurs in no other AdV genus than *Mastadenovirus*, and is highly conserved in Mastaadenoviruses ^65, 66, 67^, we speculate that the viruses lacking protein V evolved other mechanisms to secure their capsid DNA before arriving at the nucleus.

**Figure 8:**
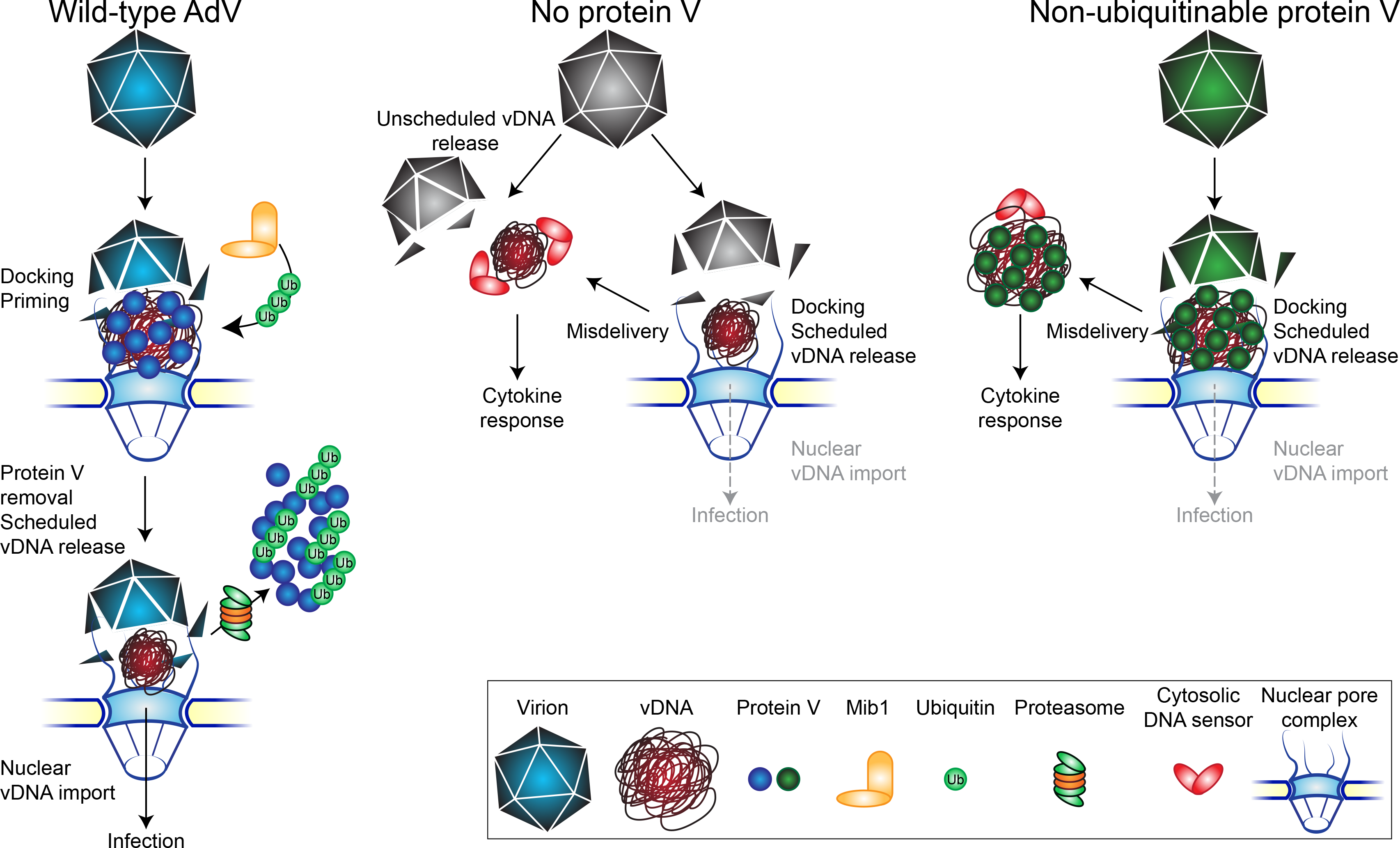
Schematic depiction of scheduled and unscheduled release of vDNA from AdV particles during virus entry into cells. Upon docking at the NPC, Mib1-mediated ubiquitination primes the AdV capsid for vDNA release and nuclear import (left). Proteasome activity removes protein V, which facilitates the import of the viral genome into the nucleus. Non-ubiquitinable protein V fails to separate from the vDNA and prevents vDNA nuclear import (right). Absence of protein V reduces the stability of the virions in the cytoplasm which increases unscheduled vDNA release distant from the NPC (center). Free cytoplasmic genomes are recognized by DNA sensors, leading to a strong cytokine response.

## DISCUSSION

Understanding how cells control virion stability is key to pathology and advances the field of synthetic virology, for example for the development of customized gene delivery vehicles and vaccines. Human AdVs and their interactions with cells are a highly advanced model of virus-host interactions at all levels, ranging from single cell infection, immunity, persistent and acute human disease to therapy and vaccination ^35, 56, 68, 69^. Human AdV particles are composed of major and minor capsid proteins conferring structural and accessory functions, such as DNA confinement, particle stability or membrane rupture ^68, 70, 71^. The latter is mediated by protein VI with an amphipathic helix interacting with sphingolipids ^72, 73, 74, 75, 76, 77, 78, 79^. Virions penetrated to the cytosol are leaky containers, and their DNA is accessible to dyes, as shown by click chemistry ^52^. They detach from nuclear envelope-proximal microtubules, bind to the NPC filament protein Nup214, and uncoat their genome through the involvement of kinesin motors and microtubules ^25, 60, 80, 81, 82^.

Our data here show how an internal linchpin protein safe-guards the virions trafficking in the cytoplasm from premature DNA release, and how the NPC-docked virions are primed for disassembly by the E3 ubiquitin ligase Mib1. The mechanism identified here is distinct from the one inactivating antibody-coated virus particles ^83^. AdVs loaded with antibodies may enter cells, for example in individuals with preexisting anti-AdV antibodies, and the cytosolic particles become subject to inactivation through recognition of the immunoglobulins by the E3 ubiquitin ligase TRIM21 and subsequent proteasomal degradation ^84^. This catastrophic disassembly process exposes vDNA in the cytosol and raises innate immunity and inflammation. The question how native AdV capsids hold their DNA and resist the cytoplasmic crowding has remained unresolved, however. This is remarkable since these non-opsonized particles have not only shed a number of stabilizing proteins, such as IIIa, VIII, VI, pentons and fibers ^50, 85^, but are also subject to pulling forces from dynein and kinesin motors during cytoplasmic transport on microtubules ^86, 87, 88, 89, 90^.

Here we show that protein V secures the viral genome in the capsid, and that its release is mediated by the E3 ligase Mib1. This process facilitates nuclear import of the viral genome. Genetic ablation of protein V reduced the efficiency of nuclear import of vDNA and thereby decreased the infectivity of the particles. However, the absence of protein V did not compromise the disassembly of the capsid or the release of the vDNA. In fact, our results show that the absence of protein V renders the particles less stable, leading to premature disassembly of the capsid before reaching the NPC. Accordingly, AdV particles lacking protein V were less thermostable than wild-type particles, possibly because about 160 copies of protein V act as a capsid glue by connecting the vDNA and protein VI ^55, 91, 92, 93^. Protein V balances the negative charges on vDNA together with proteins VII, X/μ, and IVa2, akin to a histone H1-like protein ^94^. During entry, protein V quantitatively dissociates from the particles at the NPC ^51^. This result and the observation that salt-treated vDNA cores isolated from purified virions release protein V ^95^ suggest that protein V binding to the vDNA core can be tuned.

We identified the cell tuner of protein V, the E3 ligase Mib1, and showed that the efficiency of nuclear translocation of the AdV genome depends on the ubiquitination of protein V. Ubiquitination analyses by affinity purification mass spectrometry identified two lysine residues on protein V that carried a di-glycine (di-G) remnant after tryptic digest. Di-G remnants on the ε-amino group of lysines derive from digestion of proteins conjugated to ubiquitin or ubiquitin-like modifiers, such as ISG15 and NEDD8, which cannot be distinguished from ubiquitin by MS ^48, 96^. It is unlikely that the di-G remnants on protein V were derived from another modification than ubiquitin, since Kim et al. reported that by our detection methodology >94% of K-ε-GG sites are ubiquitinated and only 6% due to NEDD8ylation or ISG15ylation ^47^. In fact, we believe that lysine residues additional to K178 and K188 on protein V are ubiquitinated during entry. In agreement with this notion, mass spectrometry and shotgun proteomics of purified AdV-C5 gave no evidence for ubiquitination of protein V residues (data not shown), consistent with lack of protein V ubiquitination evidence from liquid chromatography–high resolving mass spectrometry (LC-MS) of purified AdV-C2 particles ^97^, implying that ubiquitination of protein V occurs during entry and can be used as a toggle switch.

Protein V potentially provides a direct ubiquitination substrate for Mib1. Yet, not all viral genomes were released from AdV-C5-ΔV virions in the Mib1-KO cells suggesting that another factor besides protein V may also be ubiquitinated and involved in Mib1-priming of vDNA uncoating. We speculate that this factor is a cellular protein that stabilizes the viral capsid against disruption by kinesin-mediated pulling forces, which had been implicated in capsid disruption at the NPC ^25^. It is plausible that ubiquitination and proteasomal degradation of this factor occurs at the NPC since the proteasome inhibitor MG132 prevented the vDNA release from the NPC-docked capsid upon induction of Mib1 expression in Mib1-KO cells. The role of the proteasome in AdV capsid disassembly at the NPC represents a novel function and can now be studied in more detail. It extends the role of the proteasome beyond binding, internalization and trafficking of other viruses ^31, 98, 99, 100, 101^, and increases the therapeutic potential of proteasome inhibitors. Developing proteasomal inhibitors against vDNA uncoating may also be beneficial in reducing cytokine response and inflammation. Remarkably, genetic perturbations of protein V in AdV-C5-ΔV, AdV-C5-V-KR and AdV-C5-V-KRrev increased the levels of viral genomes in the cytosol as a result of unscheduled vDNA release or misdelivery. The AdV-C5-ΔV particles led to the largest overall induction of innate cytokines, strongly dependent on the presence of the cGAS sensor. We speculate that the non-ubiquinatable V-KR protein partially shields misdelivered vDNA from cellular sensors. Misdelivery of vDNA is, however, not unique to AdV. For example, more than half of human immunodeficiency virus-1 (HIV-1) reverse-transcribed genomes were found to be capsid-free in the cytosol of primary human macrophages ^102^. It remains to be explored how other viruses secure and shield their vDNA from cellular sensors, or activate cytosolic DNA sensors ^103, 104^.

## MATERIALS AND METHODS

### Cell culture and virus production

HeLa-ATCC, A549, HDF-TERT, and HER911 cells were maintained in Dulbecco’s Modified Eagle’s Medium (GIBCO) supplemented with non-essential amino acids (Thermo Fisher) and 10% fetal calf serum (FCS, GIBCO). During infection experiments, the medium was additionally supplemented with 100 U/ml penicillin and 100 μg/ml streptomycin. The cells were grown at 37 °C in a 5% CO_2_ atmosphere for no longer than 20 passages. HeLa-sgNT and HeLa-sgMib1 cells have been previously described ^32^. HeLa-sgMib1 cells expressing mScarlet-Mib1 were generated by lentiviral transduction followed by selection with 2 μg/ml puromycin. MPI-2 cells were maintained in RPMI medium (Sigma) supplemented with 10% fetal calf serum and 10 ng/ml granulocyte macrophage colony-stimulating factor (GM-CSF, Miltenyi Biotec) ^61, 62^.

All AdVs were grown in A549 cells and purified over two cesium chloride gradients as previously described ^50, 105^. AdV-C5 (wt300) has been previously described ^106^. AdV-C5-ΔIX was kindly provided by R. Hoeben (Leiden University Medical Center, The Netherlands) ^107^. AdV-C2-GFP-V was used as described ^51^. Capsid-labeled viruses were generated as described ^26, 108^. Genome-labeled AdV was produced by growing the virus in A549 cells in the presence of 2.5 μM EdC (5-ethynyl-2’-deoxycytidine, Jena Biosciences) as described ^52^. AdV-C5-ΔV, AdV-C5-V-KR, AdV-C5-V-KRrev, AdV-C5-V-K178R, AdV-C5-V-K188R, and AdV-C5-V-K178/188R were generated by recombineering from the pKSB2 bacmid which contains the entire AdV-C5 wt300 genome ^106, 109^.

### Generation of AdV mutants

In a first step, the galK cassette was amplified by PCR from the pGalK plasmid using primers that carried 45 bp homology sequences directly up- or downstream of the protein V coding region (see **Table S1**, primers GalK_f and GalK_r). The PCR product was purified by gel extraction and digested with DpnI (Promega) for 1 h to remove residual template DNA before a second round of purification. Electrocompetent E. coli SW102 cells harboring the AdV-C5 containing bacmid (pKSB2) were then electroporated with the purified PCR construct. Positive clones were verified by sequencing and underwent a second electroporation reaction. In the case of AdV-C5-ΔV, this was done with a dsDNA oligonucleotide (dV_f and dV_r, **Table S1**) consisting of the left and right homology sequences (Fig. 4A). For the AdV-C5-V-KR mutant, the recombination substrate was a synthesized modified protein V DNA sequence in which all lysine codons were replaced by arginine codons flanked the by left and right homology arms (**Table S1**; synthesized by Thermo Fisher Scientific). The resulting AdV-C5-V-KR bacmid then served as the template for constructing the AdV-C5-V-KRrev mutant. Bacmid DNA from positive clones was extracted, digested with PacI to release the AdV genome, and transfected into HER911 cells ^110^. Rescued viruses were plaque-purified, expanded, and verified by sequencing.

### Single-round infection assays

Ten thousand cells were seeded in a black 96-well imaging plate. On the following day, the virus was diluted in infection medium (DMEM supplemented with 2% FCS, non-essential amino acids, pen/strep) to reach an infection of about 40%. The culture supernatant was aspirated and 100 μl of diluted virus suspension was added to the cells. The cells were fixed with 4% paraformaldehyde (PFA) in PBS for 10-15 min at RT. Remaining PFA was quenched with 25 mM NH_4_Cl diluted in PBS for 5-10 min, followed by permeabilization with 0.5% Triton X-100 in PBS for 3-5 min. Cells were stained with rabbit anti-protein VI ^26^ or mouse anti-E1A clone M73 (Millipore, 05-599) diluted in blocking buffer (10% goat serum in PBS) for 1 h at 4 °C. After three washes of 4 min each in PBS, cells were stained with secondary antibody (goat anti-rabbit-AlexaFluor 488 or goat anti-mouse-AlexaFluor 488, Thermo Fisher) diluted in blocking buffer containing 1 μg/ml DAPI for 30 min at RT. After three more washes of 4 min in PBS, cells were imaged in a Molecular Devices high-throughput microscope (IXM-XL or IXMc) in widefield mode with a 20x objective. For quantification of infection with CellProfiler ^111^, nuclei were segmented according to the DAPI signal, and the intensity of the infection marker over the nuclear mask was measured.

### Virus particle infectivity

Virus input was normalized to the number of viral particles that bound to the cells, which was determined in a binding assay. To this end, HeLa cells grown on cover slips in a 24-well plate were incubated with virus for 1 h on ice, after which the virus inoculum was washed and away and cells were immediately fixed with 4% PFA. Bound viral particles were stained with mouse anti-Hexon 9C12 antibody ^112^ and goat anti-mouse-AlexaFluor 488. Nuclei were stained with DAPI and cell outlines with AlexaFluor 647-conjugated succinimidyl ester (Thermo Fisher). Samples were imaged using a Leica SP8 confocal laser scanning microscope (cLSM). Three-dimensional stacks were recorded and the number of particles that bound to cells were quantified in maximum projections using CellProfiler. HeLa cells in a 96-well plate were then incubated with a 1:2 dilution series of virus starting from 50 bound particles per cell. After 1 h on ice, virus inoculum was removed, and fresh medium was added. Cells were fixed at 20 hpi and stained for E1A as described above. MOI for infection assays was based on infectious particles in the paerticular assays. For example, MOI 0.5 indicated that 50% of the cells were infected at the time of fixation.

### Confocal microscopy

A Leica SP8 cLSM was used in all experiments, in which single viral particles and genomes were imaged. Imaging was performed with a 63x magnification oil objective with a numerical aperture of 1.40 and a zoom factor of 2, with a pixel size of 0.181 μm. z-stacks were captured with a step size of 0.5 μm to capture the entire cell, and the size of the pinhole was 1 Airy unit. Leica hybrid detectors (HyD) were used for each channel.

### SDS-PAGE and Western Blotting

Purified virus particles were lysed in SDS-PAGE lysis buffer (200 mM Tris pH 6.8, 10% glycerol, 5 mM EDTA, 0.02% bromophenol blue, 5% SDS, 50 mM DTT) and boiled for 5 min at 95 °C. Samples were then loaded onto a 10% SDS-PAGE gel and transferred to a PVDF membrane (Amersham). After blocking with blocking solution (5% milk powder in 20 mM Tris, 150 mM NaCl, 0.1% Tween-20, pH 7.5), the membrane was incubated with primary and secondary antibodies diluted in blocking solution at 4 °C overnight or 1 h at RT, with 4 washes of TBST in between. HRP-coupled secondary antibody was detected using the ECL reagent (GE Healthcare). Primary antibodies used in Western blotting were rabbit polyclonal anti-Hexon and rabbit polyclonal anti-protein V (both kind gifts from Ulf Pettersson). Secondary antibody used in Western blotting was goat anti-rabbit-HRP (Cell Signaling, 7074). Chemiluminescence signals were detected using an ImageQuant LAS 4000 system. Alternatively, after gel electrophoresis separated proteins were stained in the gel using Coomassie or QuickBlue protein stain (LubioScience).

### Click chemistry and vDNA analysis

Cells grown on cover slips were infected with genome-labeled AdV for various timepoints. Where specified, cells were incubated with specific inhibitors (50 nM LMB, 5 μM DBeQ, 10 nM MLN9708, 10 μM Nocodazole) 1 h prior, during and after virus inoculation at 37 °C. For microtubule disruption, cells were incubated in addition for 1 h on ice in the presence of 10 μM Nocodazole 1h before virus inoculation. After fixation, quenching, and permeabilization, samples were stained for incoming capsids with the 9C12 anti-hexon antibody. After primary and secondary antibody incubation, the cover slips were inverted onto a 30 μl droplet of click reaction mix for 2 h at RT. The freshly prepared click reaction mix consisted of 10 μM AlexaFluor 488-conjugated azide (Thermo Fisher Scientific), 1 mM CuSO4, and 10 mM sodium ascorbate in the presence of 1 mM THPTA (Sigma) and 10 mM aminoguanidine (Sigma) in PBS. Samples were stained with DAPI and AlexaFluor 647-conjugated succinimidyl ester and imaged with a Leica SP8 cLSM as described above. Nuclei and single viral genomes and/or capsids were segmented according to the corresponding signal using CellProfiler. Genomes were classified as capsid-positive based on their corresponding Hexon signal. Nuclear genomes were those that overlapped with the nuclear mask that was created based on the DAPI signal. Percentage of nuclear genomes is set in relation to all capsid-free genomes.

### Confocal spinning-disk live microscopy

Eight thousand HeLa-sgMIB1 cells expressing mScarlet-MIB1 were seeded in a 10-well CELLview slide (Greiner Bio-One) with a 175 μm thick cover glass embedded in its bottom. After two days, the cells were incubated with AdV-C2-GFP-V-atto647 ^51^ at 37 °C for 30 min. Unbound virus was washed away, fresh medium without phenol-red was added to the cells, and live imaging was started on a Visitron CSU-W1 spinning disk microscope consisting of a Nikon Eclipse T1 microscope and a Yokogawa confocal scanning unit W1 with a stage top incubation system at 37 °C and 5% CO2. Z-stacks consisting of four steps with a step size of 1.4 μm were acquired every 10 s for up to 30 min with a 100x oil objective (NA 1.4) and a pinhole of 50 μm. The focus was maintained with a perfect focus system (PFS).

### Di-glycine immunoprecipitation and mass spectrometry analysis

Di-glycine immunoprecipitation (IP) and MS analysis was performed essentially as described ^113^. HeLa-sgNT cells were seeded in three 15 cm dishes to a confluency of ca. 90%. Cells were pretreated for 2 h with 10 μM MG132 (Sigma Aldrich) before addition of 250 μg of AdV-C5. After incubation at 37 °C for 2 h, cells were washed two times with PBS and trypsinized. Cells were pelleted by centrifugation at 500 xg at 4 °C for 4 min, resuspended in PBS, and centrifuged again. The cell pellet was resuspended in 1.3 ml of ice-cold lysis buffer (8 M urea, 50 mM Tris (pH 8), 150 mM NaCl, 1 mM EDTA, 50 μM PR-619, protease inhibitor cocktail (Roche)) and sonicated three times for 1 min with 1 min on ice in between using a Hielscher sonicator. Samples were then centrifuged at 16,100 xg for 15 min at 4 °C, and the supernatant was collected. Protein concentration was determined via micro BCA assay (Thermo Fisher). Five mg of the lysate was reduced and alkylated by incubation with 10 μM TCEP and 40 μM chloroacetamide for 30 min at RT in the dark. The lysate was subsequently diluted to a concentration of 4 M urea by adding 50 mM Tris (pH 8) and digested with 20 μg of LysC (Wako Chem) for 4 h at 37 °C under agitation. The lysate was then diluted to <2 M urea and incubated with 50 μg trypsin (Promega) overnight at 37 °C under agitation. The following day, 1% trifluoroacetic acid (TFA) was added to the sample followed by incubation on ice for 15 min. After centrifugation for 10 min at 3000 xg, the supernatant was transferred to a new tube and desalted using a 500 mg tC18 Sep-Pak cartridge (Waters) with a vacuum manifold according to the manufacturer’s instructions. Peptides were eluted with a 60% acetonitrile (ACN)/0.1% TFA solution and dried in a SpeedVac vacuum concentrator (Thermo Fisher). For the IP of di-glycine peptides, the sample was incubated with crosslinked agarose-coupled PTMScan Ubiquitin Remnant Motif antibody (Cell Signaling Technology) according to the manufacturer’s instructions. Following elution with 0.15% TFA, samples were lyophilized and fractionated using C18 disks at high pH through elution with increasing concentrations of ACN (2.5%, 7.5%, 12.5%, and 50% in 25 mM ammonium formate, pH 10). Eluted samples were lyophilized in a SpeedVac and stored at −20 °C until MS analysis. Samples were reconstituted in 3% ACN/0.1% formic acid (FA) for analysis by LC-MS/MS on an Orbitrap Q Exactive HF mass spectrometer (Thermo Fisher) coupled to a nano EasyLC 1000 (Thermo Fisher). For this, peptides were loaded onto a reverse-phase C18 (ReproSil-Pur 120 C18-AQ, 1.9 mm, Dr. Maisch GmbH) packed self-made column (75 mm3, 150 mm) that was connected to an empty Picotip emitter (New Objective). Solvent compositions in channels A and B were 0.1% FA in H_2_O and 0.1% FA in ACN, respectively. Peptides were injected into the MS at a flow rate of 300 nl/min and were separated using a 120 min gradient of 2% to 35% buffer B. The MS was set to acquire full-scan MS spectra (300-1700 m/z) at a resolution of 60,000. A top 12 method was used for data-dependent acquisition (DDA) mode.

Raw files were analyzed using the Proteome Discoverer software, v2.1 (Thermo Fisher). Parent ion and tandem mass spectra were searched against the UniProtKB Homo sapiens and Human Adenovirus C5 databases using the SEQUEST algorithm. For the search, the enzyme specificity was set to trypsin with a maximum of two missed cleavage sites. The precursor mass tolerance was set to 10 ppm and the fragment mass tolerance to 0.02 Da. Carbamidomethylation of cysteines was set as a fixed modification; N-terminal acetylation, oxidation of methionine, and di-glycine lysines were searched as dynamic modifications. The datasets were filtered on posterior error probability to achieve 1% false discovery rate on protein and peptide level.

### Electron microscopy

Graphite coated electron microscopy grids were treated with five microliters of glycerol free purified virus for five minutes. Grids were washed three times with distillated water and stained for thirty seconds with ten microliters of a 2% uranyl acetate solution. Samples were imaged in a FEI CM100 electron microscope at 80 keV.

### SLO penetration assay

Streptolysin O (SLO) penetration assay was essentially performed as described ^114^. Eighty thousand A549 cells were seeded onto coverslips. On the following day, cells were incubated with ca. 50 bound virus particles of capsid labeled viruses (atto565 or AlexaFluor 488) on ice for 1 h. Inocula were washed away with cold binding medium and cells were incubated at 37 °C in a water bath. SLO binding buffer (Hepes 7.4 25 mM, KOAc 110 mM, MgOAC 2.5 mM, CaCl2 0.2 mM, EGTA 1 mM, DTT 1 mM) was supplemented with 1 μg of SLO and left at RT for 5 min for activation. Cells were incubated in SLO binding buffer with or without SLO for 10 min on ice. Cells were washed twice with SLO binding buffer and incubated for 5 min at 37 °C to allow polymerization. Cells were washed once with SLO Binding buffer and twice with SLO internalization buffer (Hepes 7.4 25 mM, KOAc 110 mM, MgOAC 2.5 mM, EGTA 2 mM). Cells were incubated with mouse anti-Hexon 9C12 or rabbit anti-AlexaFluor 488 (Molecular Probes) in SLO internalization buffer for 1 h on ice. Samples were washed three times with SLO internalization buffer and fixed with 3% PFA in 25 mM Hepes-KOH pH 7.4, 110 mM KOAc, 2.5 mM MgOAc for 15 min at RT. Samples were quenched and stained with secondary antibody goat anti-mouse-AlexaFluor 488 or goat anti-rabbit-AlexaFluor 596 (Thermo Fisher) diluted in blocking buffer containing DAPI for 1 h at RT. Cells were washed, treated with succinimidyl ester AlexaFluor 647, and mounted as described above. A Triton X-100 permeabilized sample was included to control the accessibility of the antibodies. Samples were imaged with a Leica SP8 upright confocal microscope. For image analysis, virus particles were segmented according to their capsid label signal and classified as cytosolic based on their antibody staining signal.

### Intracellular trafficking of viral particles

Eighty thousand A549 cells were seeded on coverslips. On the following day, cells were incubated with ca. 80 bound particles of each virus on ice as described above. Inocula were removed and cells were incubated with infection medium at 37 °C. Cells were fixed, quenched and permeabilized. Samples were stained with mouse anti-Hexon 9C12 and goat anti-mouse AlexaFluor 488 in blocking buffer containing DAPI. Cells were treated with succinimidyl ester AlexaFluor 647 and mounted onto glass holders. Samples were imaged with a Leica SP8 upright confocal microscope. For image analysis, virus particles were segmented according to their Hexon signal and masked with the nuclear signal.

### Thermostability assays

Two micrograms of purified virus particles were incubated in a thermocycler for 5 min at 35, 40, 43, 47, 50, 55 or 60 °C. Samples were cooled down on ice and incubated with 5 μM DiYO-1 (AAT Bioquest) for 5 min at RT. Fluorescence was recorded with a Tecan Infinite M200 plate reader (excitation 490 nm, emission 520 nm). To assess the influence of heat treatment on infection, 10,000 A549 cells were seeded in a black 96-well imaging plate. On the following day, virus suspensions were heat treated for 5 min at 40, 40.3, 40.9, 42, 43.4, 44.8, 46.2, 47.6, 49, 50 or 80 °C. Samples were placed in infection medium and added to the cells. After 24 h, cells were fixed, quenched and permeabilized. Cells were stained with rabbit anti-protein VI and goat anti-rabbit AlexaFluor 488 in blocking buffer containing DAPI. Cells were imaged in an IXMc high-throughput microscope, and infection was quantified based on the protein VI signal over segmented nuclei.

### Cytokine induction assay

Virus input was normalized to the number of viral particles that bound to MPI-2 cells, which was determined in a binding assay as described above. Four hundred and fifty thousand MPI-2 wt, shcGAS, or shEGFP cells were seeded in a 24-well plate. On the following day, the virus was diluted in cold binding medium to reach 350 bound virus particles and added to the cells for 30 min at 37 °C. Inocula were removed, cells were washed two times with cold binding medium, and fresh growth medium was added for a total of 5 or 10 h of infection. Supernatant (SN) was collected, cells were treated with 200 μl of PBS/1.5 mM EDTA for 3 min and collected along the SN. Extracts were centrifuged at 250 xg for 5 min, SN were stored at −80°C and cell pellets were re-suspended in 300 μl of TRIzol reagent (Thermo Fisher). Total RNA was extracted using Direct-zol RNA kit (Zymo Research) and the concentration was measured in a nanodrop. Two hundred nanograms of purified RNA were reverse-transcribed using Moloney Murine Leukemia Virus Reverse Transcriptase (MMLV RT, Promega). RT was primed with oligo dT(15) (Promega) and held for 1 h at 42 °C, followed by a denaturation step for 10 min at 95 °C. Samples amplified in the absence of RT were used to control for the presence of genomic DNA. One microliter of the synthesized cDNA was mixed with 0.5 μl of 10 μM pre-mixed primer and SYBR Green JumpStart (Sigma) to reach a final volume of 10 μl, and loaded in a MicroAmp Optical 96-well Plate (Applied Biosystems). Forty cycles of amplification at 60 °C were conducted in a QuantStudio 3 Real-Time PCR System (Applied Biosystems). Amplification curves were analyzed using QuantStudio Design & Analysis Software v1.5.1. Relative levels of expression compared to a non-infected control were calculated through the 2^(ΔΔCt) method ^115^, using HPRT cDNA as an endogenous loading control. Supernatants collected from these experiments were used for the chemokine and IFNβ production quantifications. CCL2 and CCL5 protein levels were quantified using a sandwich ELISA procedure (Sigma, Cat# RAB0055 & RAB0077) following the manufacturer’s instructions. Absorbance at 450 nm was measured in a Tecan Infinite M200 plate reader.

### IFNβ measurements

Thirty thousand MEF-Mx2-Luc-βKO cells were seeded per well in a 96-well plate ^62^. On the following day, MPI-2 derived supernatants were diluted 1/10 and 1/100 in fresh media and added to the cells for 20 hours. In parallel, recombinant mouse IFNβ (kindly provided by Peter Staeheli) was serially diluted and used as a standard to assess IFNβ Units per ml. Twenty hours post inoculation SN were discarded, cells were washed once with PBS and lysed in 40μl of 1 x CCLR buffer (Promega) for 10 min on a rocking plate at RT. Twenty five microliters from each lysate were transferred to a white Nunclon 96-well plate (ThermoFisher). Thirty microliters of Luciferase Assay Reagent (LAR, Promega) were added to the wells using a TECAN plate reader with injection unit, followed by shaking for 2 seconds and integration of the luminescence signal over 10 seconds.

### RNA FISH with branched DNA signal amplification assay

Forty-five thousand MPI-2 wt cells were seeded in a 96-well plate. On the following day, the virus was diluted in cold binding medium to reach 350 bound virus particles and added to the cells for 30 min. Five or ten hours post infection cells were fixed with 3% PFA in PBS for 30 min at RT, washed twice with PBS, and dehydrated by subsequent incubation with 50%, 70% and 100% ethanol for 2 min at RT. Dehydrated samples were stored at −20 °C until staining. Samples were rehydrated by incubation with 70%, 50% ethanol and PBS for 2 min at RT. Rehydrated samples were FISH-stained against mouse Ccl2 mRNA using ViewRNA mRNA FISH assay (type 1 probe, Alexa Fluor 546, #6006661-01, probes were made against the sequence between mouse Ccl2 gene map positions 2-785, ThermoFisher) according to the manufacturer’s instructions. Subsequently, cells were incubated in PBS containing DAPI and succinimidyl ester AlexaFluor 647 (Thermo Fisher) for 10 min at RT. Cells were imaged in an ImageXpress Micro confocal miroscope (Molecular Devices) (60 μm pinhole, 15 stacks, 1.5 μm slice thickness) with a 40x objective. For quantification, cells were segmented according to the DAPI and succinimidyl ester signals, and segmented Ccl2 dots were then related to the cells using CellProfiller ^111^.

### Statistical analysis

All graphs display mean ± standard deviation (SD) unless stated otherwise. Statistical tests used are indicated in the figure legends. ns, not significant; *, p<0.05; **, p<0.01; ***, p<0.001).

**Table S1.**
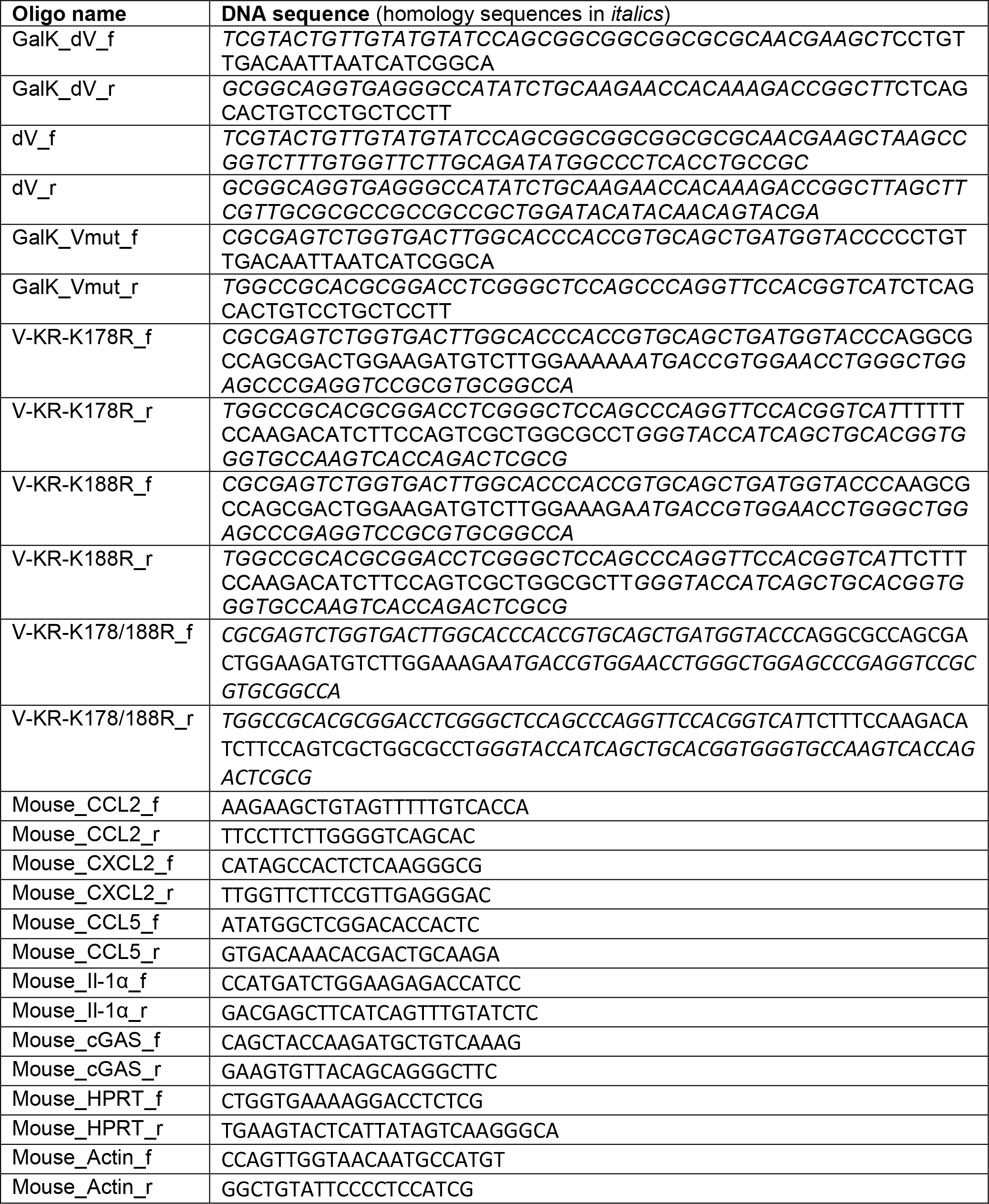
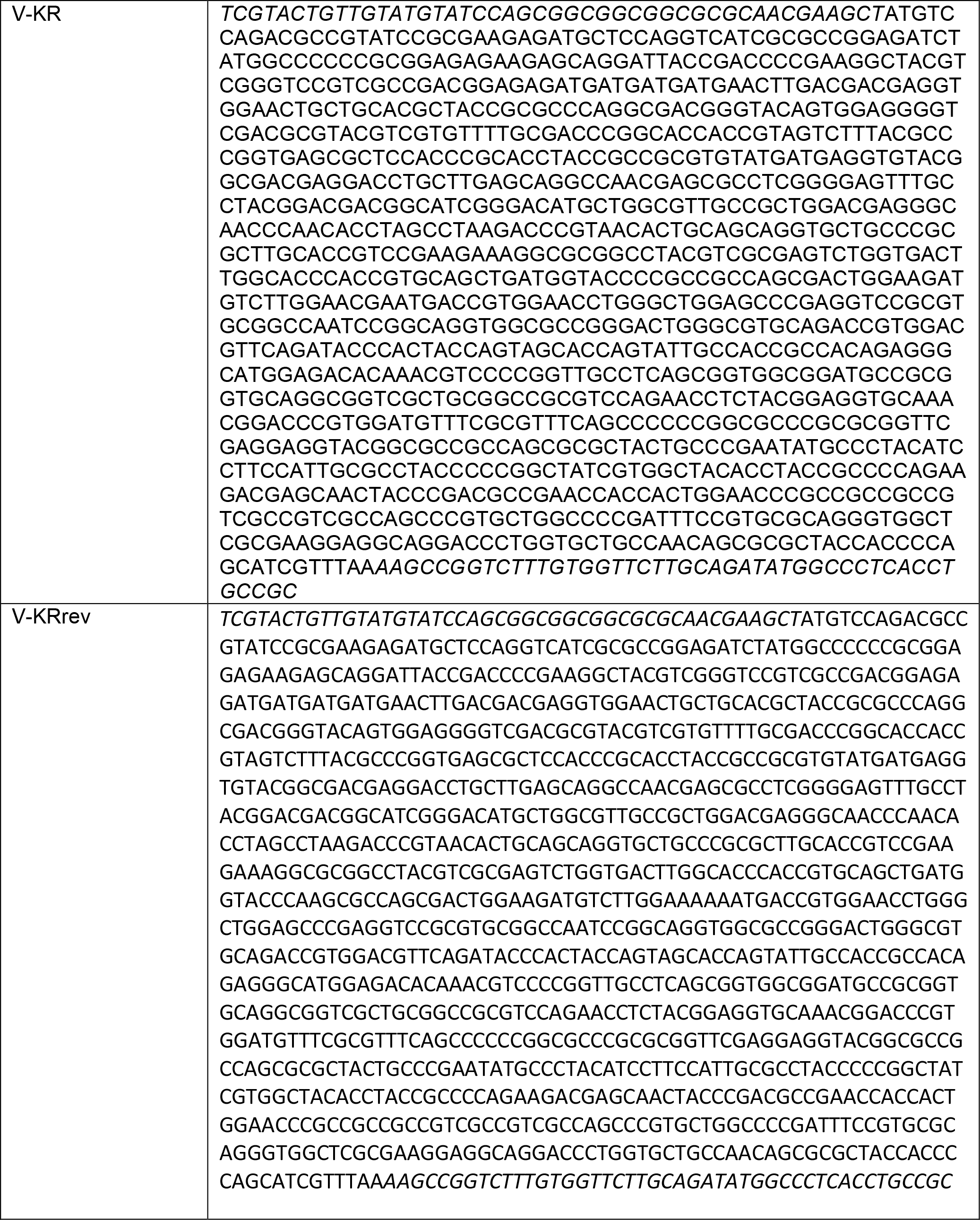

## Supporting information

Variant calling Ad5-dV

## SUPPLEMENTAL FIGURES

**Figure S1:**
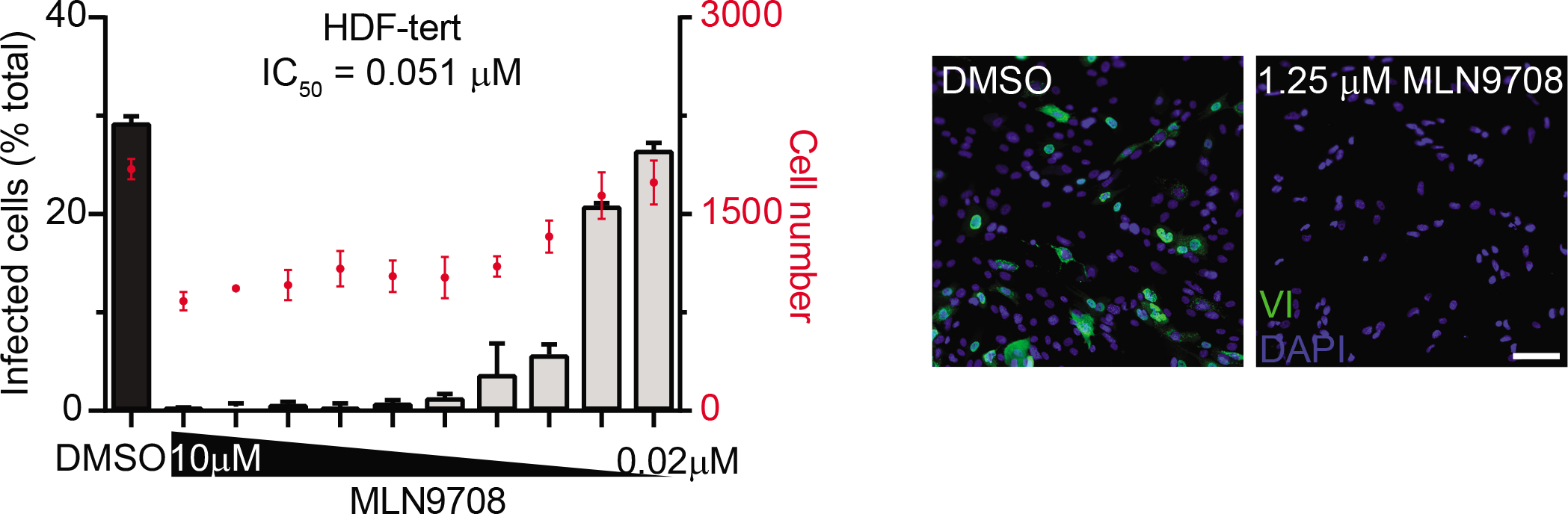
Proteasome inhibition reduces AdV infection in non-cancer cells. HDF-TERT cells were infected with AdV-C5 at a MOI of 0.3 for 24 h in the presence of DMSO or varying concentrations of proteasome inhibitor MLN9708. Representative images are shown. After fixation, cells were stained with anti-pVI antibody and DAPI. Infection was scored by percentage of pVI-positive nuclei. Scale bar, 100 μm.

**Figure S2:**
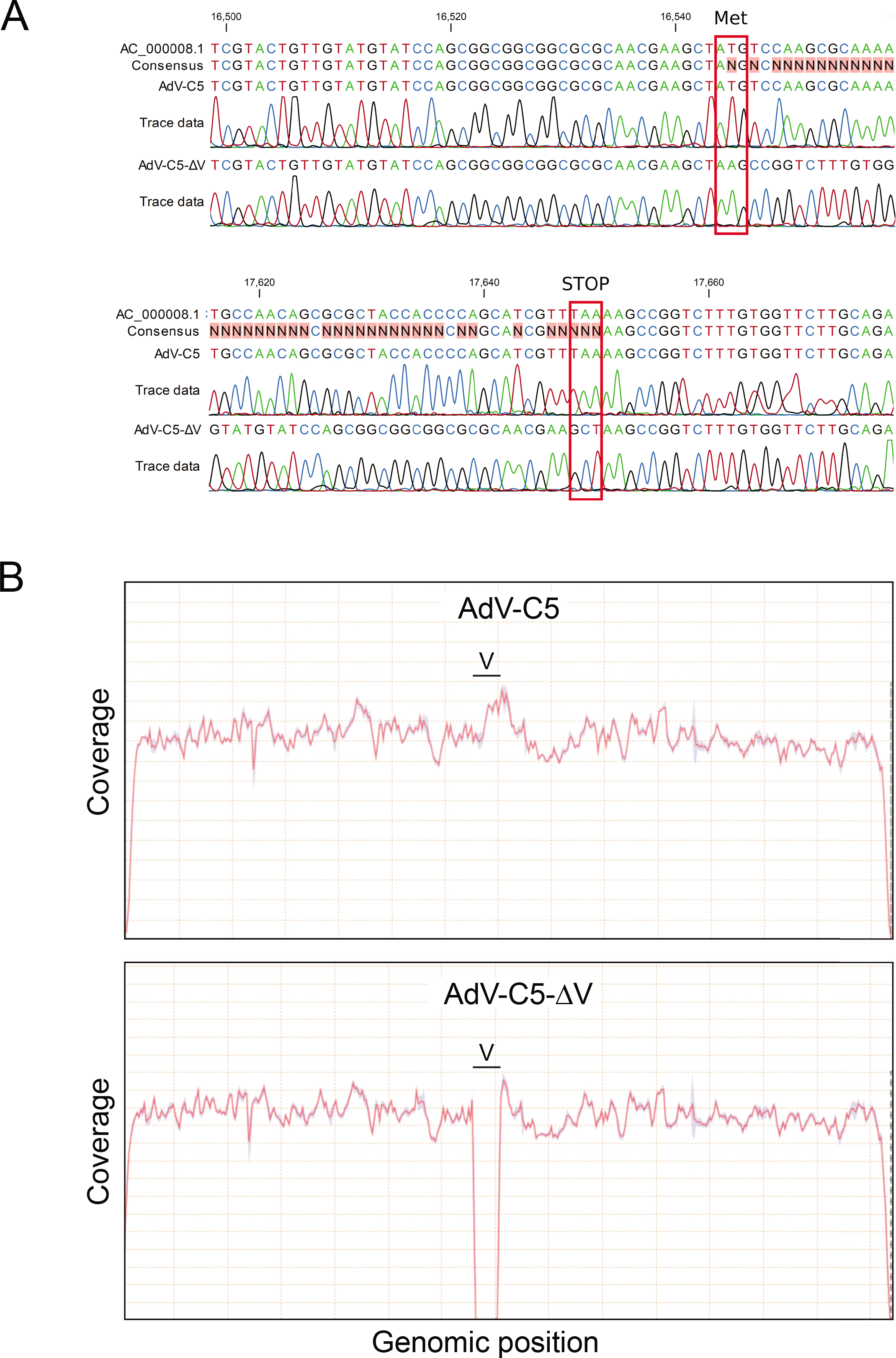
Sequencing analysis of AdV-C5-ΔV confirms deletion of protein V coding region. (A) Protein V genomic region was PCR amplified from purified AdV-C5 and AdV-C5-ΔV viral genomes. Band purified PCR amplicons were sequenced using two internal primers. Sequences were aligned to wild-type AC_000008.1 reference. Non-matching nucleotides are highlighted with red in the consensus sequence. (B) Reads obtained from a short read Illumina NGS run were mapped to AC_000008.1 reference using a Bowtie2 algorithm. Nucleotide coverage is displayed in function of AC_000008.1 genomic position for both AdV-C5 and AdV-C5-ΔV.

**Figure S3:**
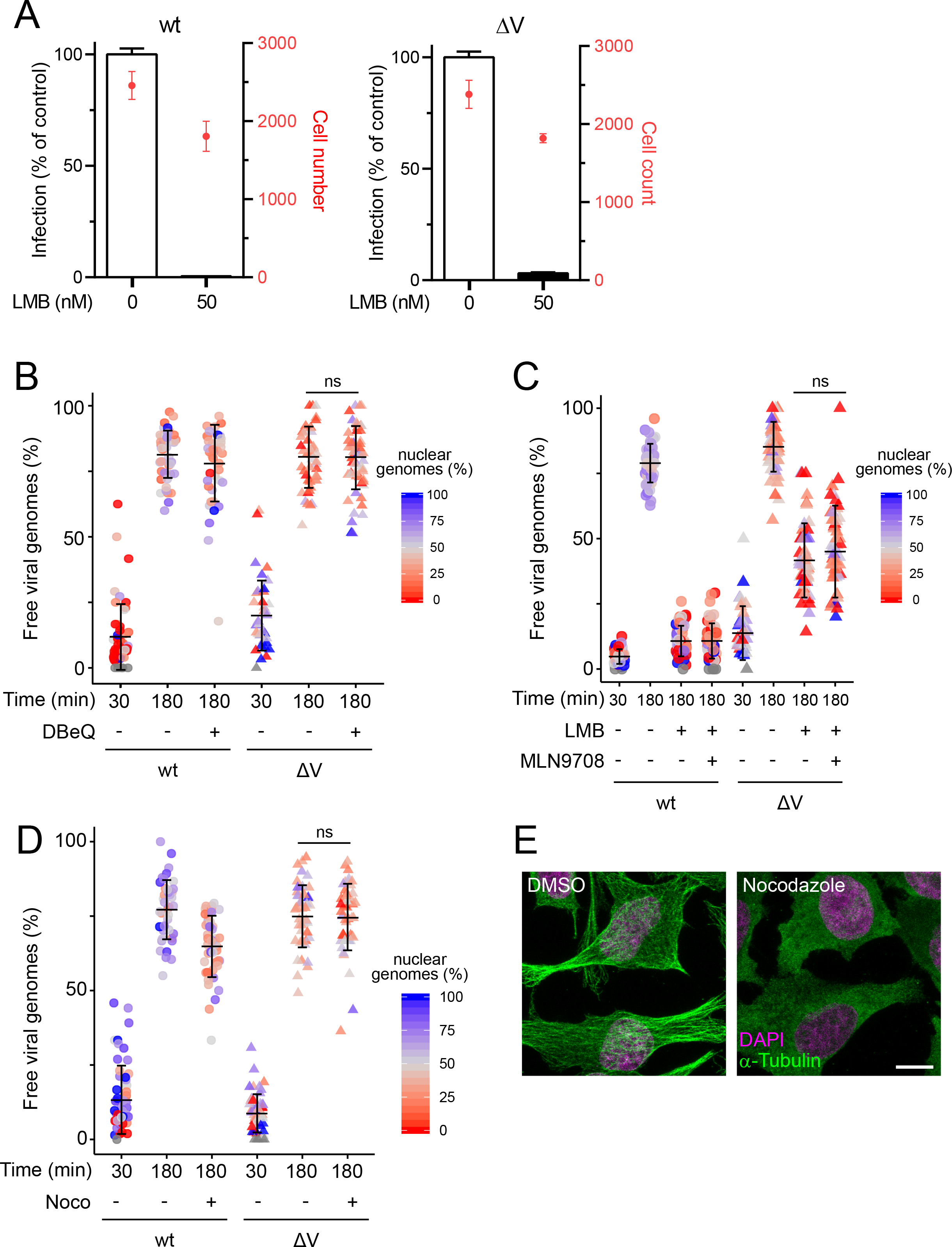
Unscheduled genome release from AdV-C5-ΔV is independent of p97, the proteasome, and microtubules. (A) HeLa cells were infected with AdV-C5 or AdV-C5-ΔV at a MOI of 0.5 for 24 h in the presence of vehicle or 50 nM LMB. After fixation, cells were stained with anti-pVI antibody and DAPI. Infection was scored by percentage of pVI-positive nuclei. Graphs show the mean ± SD. (B) HeLa cells were incubated with genome-labeled AdV-C5 or AdV-C5-ΔV for 30 or 180 min in the absence or presence of 5 μM DBeQ. Cells were fixed and stained with anti-Hexon and vDNA using click chemistry. Data are shown as mean ± SD. Ratio of the number of capsid-free genomes over the nuclear mask has been color-coded. Statistical significance was assessed by using a nonparametric ANOVA (Kruskal-Wallis test) with Dunn’s correction for multiple comparisons. ns, not significant. (C) HeLa cells were incubated with genome-labeled AdV-C5 or AdV-C5-ΔV for 30 or 180 min in the absence or presence of 50 nM LMB and/or 10 μM MLN9708. Cells were fixed and stained with anti-Hexon and vDNA using click chemistry. Data are shown as mean ± SD. Ratio of the number of capsid-free genomes over the nuclear mask has been color-coded. Statistical significance was assessed by using a non-parametric ANOVA (Kruskal-Wallis test) with Dunn’s correction for multiple comparisons. ns, not significant. (D, E) HeLa cells were incubated with genome-labeled AdV-C5 or AdV-C5-ΔV in the absence or presence of 10 nM Nocodazole. Data are shown as mean ± SD. Ratio of the number of capsid-free genomes over the nuclear mask has been color-coded. Statistical significance was assessed by using a non-parametric ANOVA (Kruskal-Wallis test) with Dunn’s correction for multiple comparisons. ns, not significant. Tubulin staining shows depolymerization of microtubules. Images are maximum projections. Scale bar, 10 μm.

**Figure S4:**
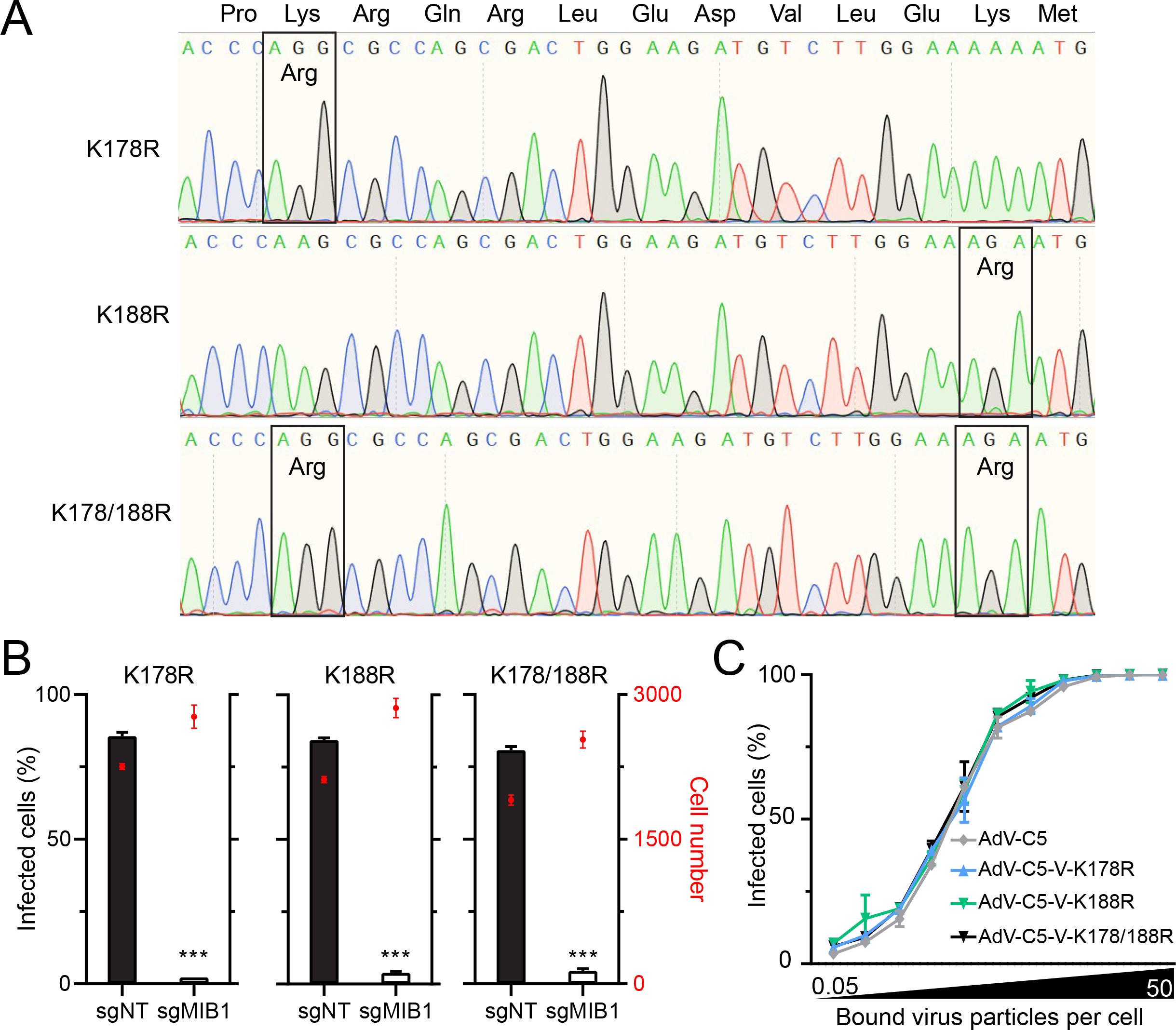
Mutations at protein V K178 and K188 have no effect on particle infectivity and dependence on Mib1. (A) Spectra from Sanger sequencing of DNA extracted from AdV-C5-V-K178R, AdV-C5-V-K188R, and AdV-C5-V-K178/188R viruses. (B) HeLa-sgNT and sgMib1 cells were infected with AdV-C5-V-K178R, AdV-C5-V-K188R, or AdV-C5-V-K178/188R for 24 h. Cells were fixed and stained with anti-protein VI and DAPI. Infection was scored by percentage of pVI-positive nuclei. Data are shown as mean ± SD. Statistical significance was assessed using unpaired T-tests. ***, p<0.001. (C) HeLa cells were incubated with the indicated viruses at an input normalized to the number of bound viral particles. After 1 h on ice, virus inoculum was removed and fresh medium was added. Cells were fixed at 24 hpi and stained for E1A and DAPI. Infection was scored by percentage of E1A-positive nuclei. Data are shown as mean ± SD.

**Figure S5:**
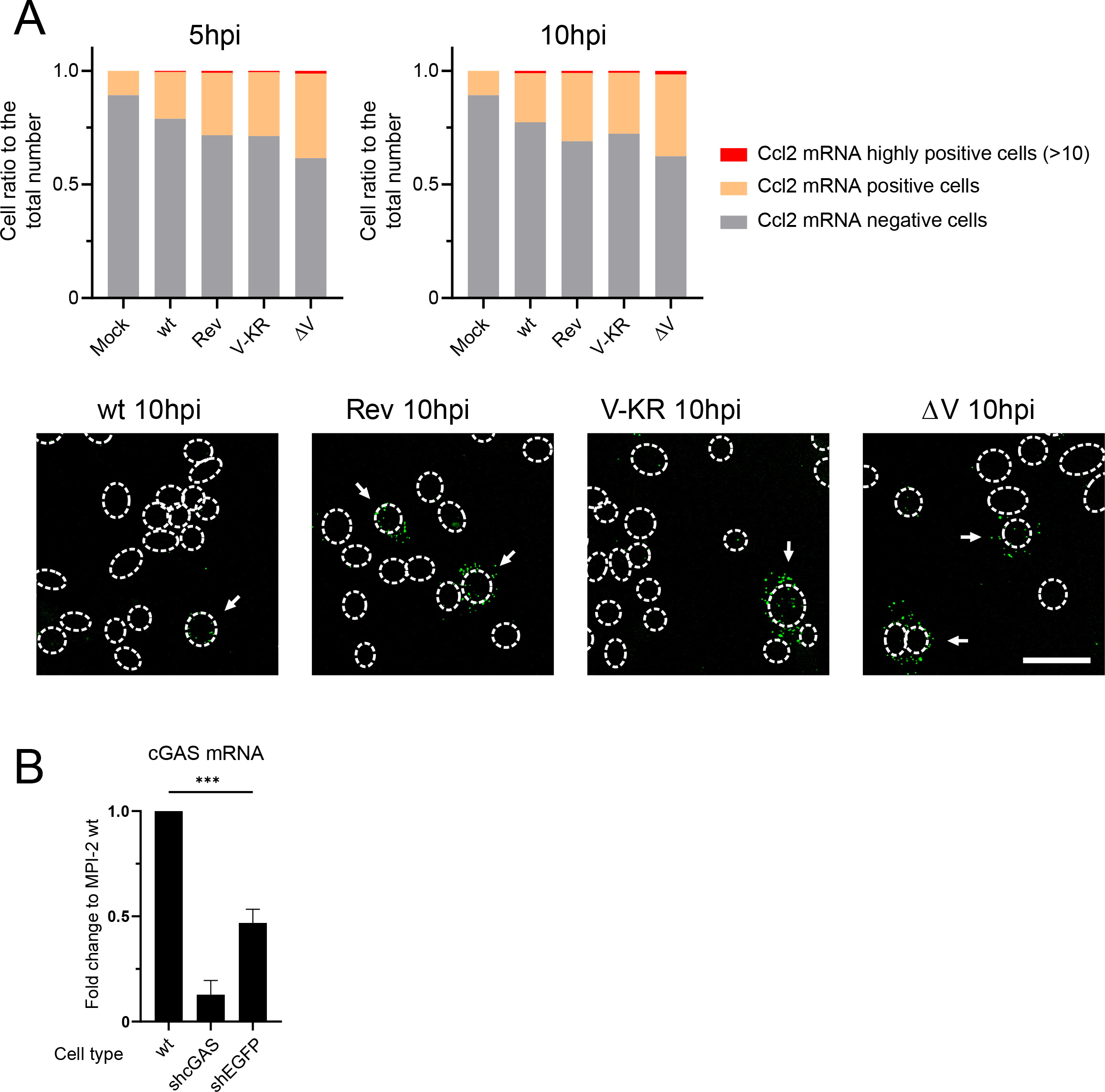
Protein V mutants induce stronger cytokine response than wild-type virus. (A) MPI-2 cells were infected with AdV-C5, AdV-C5-VKRrev, AdV-C5-V-KR, or AdV-C5-ΔV for 5 or 10 h. Cells were fixed and stained against Ccl2 mRNA using RNA FISH with branched DNA signal amplification. Nuclei were stained with DAPI and cell outlines with succinimidyl ester. Ccl2 mRNA dots were segmented and quantified per cell. Images are maximum projections. Nuclear outlines are based on DAPI. Arrowheads indicate high Ccl2 expressing cells. Scale bar, 20 μm. (B) MPI-2 wt, shcGAS, or shEGFP cells were lysed and processed for RT-qPCR. Fold change in cGAS mRNA expression was addressed through the 2^-(ΔΔCt) method using HPRT as an endogenous control. ***, p<0.001.

**Movie S1.** HeLa-sgMib1 cells expressing mScarlet-Mib1 were incubated with AdV-C2-GFP-V-atto647 at 37°C for 30 min. The virus inoculum was removed, and cells were placed in a confocal spinning-disk microscope. Timestamps are in min:sec. Arrows indicate GFP-V dissociation events. Three capsids containing GFP-V are visible at the nucleus, of which the two particles on the left discharge their GFP-V. Scale bar, 10 μm.

## ACKNOWLEDGMENTS

We thank Leta Fuchs, Nicole Meili, and Melanie Grove for help with generating the protein V mutants and electron microscopy, and Tobias Kockmann and the Functional Genomics Center Zurich for help with mass spectrometry. We thank Yohei Yamauchi and all members of the Greber lab for helpful discussions. The work was supported by grants from the Swiss National Science Foundation to UFG (31003A_179256 / 1; and 2014/264).

## AUTHOR CONTRIBUTIONS

**Conceptualization:** Michael Bauer, Urs F. Greber

**Data curation:** Michael Bauer, Alfonso Gomez-Gonzalez

**Formal analysis:** Michael Bauer, Alfonso Gomez-Gonzalez

**Funding acquisition:** Urs F. Greber

**Investigation:** Michael Bauer, Alfonso Gomez-Gonzalez

**Methodology:** Michael Bauer, Alfonso Gomez-Gonzalez, Maarit Suomalainen

**Project administration:** Urs F. Greber

**Resources:** Silvio Hemmi, Maarit Suomalainen

**Software:** Michael Bauer, Alfonso Gomez-Gonzalez

**Supervision:** Urs F. Greber

**Validation:** Michael Bauer, Alfonso Gomez-Gonzalez

**Visualization:** Michael Bauer, Alfonso Gomez-Gonzalez

**Writing & editing:** Michael Bauer, Alfonso Gomez-Gonzalez, Maarit Suomalainen

**Final text:** Urs F. Greber with input from all coauthors

